# CryoEM reveals the structure of an archaeal pilus involved in twitching motility

**DOI:** 10.1101/2023.08.07.552258

**Authors:** Matthew C. Gaines, Shamphavi Sivabalasarma, Michail N. Isupov, Risat Ul Haque, Mathew McLaren, Cyril Hanus, Vicki A.M. Gold, Sonja-Verena Albers, Bertram Daum

## Abstract

Amongst the major archaeal filament types, several have been shown to closely resemble bacterial homologues of the Type IV pili (T4P). Within *Sulfolobales,* member species encode for three types of T4P, namely the archaellum, the UV-inducible pilus (Uvp) and the archaeal adhesive pilus (Aap). Whereas the archaellum functions primarily in swimming motility, and the Uvp in UV-induced cell aggregation and DNA-exchange, the Aap plays an important role in adhesion and twitching motility. All previously solved Aap appear to have almost identical helical structures. Here, we present a cryoEM structure of the Aap of the archaeal model organism *Sulfolobus acidocaldarius.* We identify the component subunit as AapB and find that while its structure follows the canonical T4P blueprint, it adopts three distinct conformations within the pilus. The tri-conformer Aap structure that we describe challenges our current understanding of pilus structure and sheds new light on the principles of twitching motility.

## Introduction

Archaea possess a large variety of filamentous cell surface extensions, which function in adhesion to (a)biotic surfaces, biofilm formation, DNA exchange, cell-cell recognition, exchange of nutrients and motility in liquid environments ^1^. These include ABP filaments from *Pyrobaculum calidifontis* ^2^ and the *Sulfolobus acidocaldarius* threads, which structurally resemble bacterial type-I pili, but which are likely assembled by a distinct mechanism ^3^. Furthermore, *Methanothermobacter thermoautotrophicus* generates fimbriae, which are important for biofilm formation ^4^, and a structure of the filamentous archaeal DNA import machinery related to conjugative DNA transfer systems has also been very recently reported ^5^. Finally, archaea-specific filaments exist, such as unusual, barbed wire like Hami in *Altarchaeum hamiconexum* ^6^ and canulae in *Pyrobaculum calidifontis* ^2^, which both appear to have important roles in adhesion to surfaces and other cells. However, perhaps the so-far best characterised are those homologous to T4P, which are common in archaea and bacteria.

Many archaeal genomes contain operons encoding T4P ^7^. Archaeal T4P assembly are generally thought to be simpler than those found in bacteria, and mainly contain an ATPase, an inner membrane platform protein, and a set of pilin proteins. The latter exhibit a canonical class III signal peptide that is processed via a PibD protein prior to their assembly into the pilus fibre ^8^. The best investigated archaeal T4P is the archaellum, which acts as a gyrating filamentous propeller that enables cells to swim through liquid media ^9–11^. As is typical for T4P, the archaellum has a helical symmetry and is highly glycosylated ^12–19^. Six structures of archaella have been solved by CryoEM to date ^14, 16–18, 20^. These revealed that archaellins follow the structural blueprint for T4P, including the conserved hydrophobic α-helix and the more variable globular C-terminal domain. The N-terminal α-helices bundle to form the core of the filament, while the C-terminal globular domains face outside ^14, 16, 17^. The archaellum can consist of one repeating subunit, or have complex heteropolymeric composition, as revealed by a recent structure from *Methanococcus villosus* ^18^.

Non-rotary archaeal T4P are thought to function as adhesives to various biotic and abiotic surfaces, and to enable DNA exchange and intercellular communication ^1, 21^. Furthermore, archaeal T4P have been shown to serve as receptors for a range of archaeal viruses ^22^. Hyperthermophilic strains such as *Sulfolobus islandicus* are infected by the rod-shaped viruses 2 and 8 (SIRV2 and SIRV8, respectively; family *Rudiviridae*) ^23, 24^ and others, such as *Saccharolobus solfataricus* by the turreted icosahedral virus (STIV; family *Turriviridae*) ^22, 23^. Another well-studied pilus is the Uvp, which is formed by various *Sulfolobus* species in response to DNA damage by e.g. UV light and leads to species specific cell-cell aggregation ^25, 26^. This allows the cells to exchange chromosomal DNA for homologous recombination ^27, 28^, likely using a bacterial-like DNA-exchange apparatus.

In *S. acidocaldarius*, the Aap has previously been described as an adhesive filament that is important for biofilm formation, and low resolution cryoEM map has been published ^29^. Based on this map, it was for the first time suggested that indeed, archaeal pili are homologous to bacterial T4P ^29^. These pili are found in all *Sulfolobales* genomes and a comparison of known homologous adhesive pili from closely related organisms such as *S. islandicus* or *Saccharolobus solfataricus* shows that these filaments have a high degree of glycosylation, possibly as a protective adaption to their harsh natural environments ^30, 31^.

New interest in Aap was triggered through the findings of a recent study, showing that this pilus is involved in archaeal twitching motility ^32^. Twitching motion is a coordinated process that begins with the extension of T4P from the bacterial cell surface. The pili then attach to a substrate, such as a solid surface or another bacterial cell, and retract, pulling the bacterial cell towards the attachment site. This process is repeated, resulting in a series of jerky movements that propel the bacteria across surfaces ^33^. Twitching motility is particularly well studied in Gram negative bacteria, including pathogenic and environmental species, and plays crucial roles in bacterial pathogenesis, biofilm formation, and nutrient acquisition ^34–39^. In such species, this process is often driven by a complex molecular machine, which includes motor ATPases that provide the energy for pilus extension (PilB) ^40^, and retraction (PilT) ^34, 41^, a cell- membrane integral platform protein (PilC) ^41^, an outer membrane-spanning conduit (PilQ) ^42^, and accessory proteins forming a periplasmic cage-like structure, called “alignment complex”, composed of PilM, N, O and P ^41^. While archaeal genomes encode for homologs of the platform protein PilC and the assembly ATPase PilB, genes for retraction ATPases akin PilT, PilQ or the PilMNOP complex are not found in archaeal T4P assembly clusters ^7^. Two pilins called *aapA* and *aapB* are present in the *S. acidocaldarius aap* pilus cluster. However, so far it was not clear whether both pilins form the pilus, as it is the case for the archaellum of *M. villosus* ^18^, or if the filament is mainly composed of one of the two gene products.

T4P are comprised of thousands of copies of the major pilin subunit(s), plus low-abundance minor pilins ^43^. Major pilins make up the bulk of the filament, while minor pilins are thought to form cell-proximal or cell-distal cap structures and are often essential for the assembly of the pilus. Both, the major and minor pilins adopt a tadpole-like structure consisting of an N-terminal ⍺-helix (α1), followed by a globular β-strand-rich C-terminal head domain ^44^. Here, we determined the structure of the Aap, and show that it is solely composed of multiple copies of AapB. Strikingly, instead using a second pilin, AapB adopts three distinct conformations within the pilus. This gives the Aap a unique, tri-conformer architecture, which has previously not been seen in any bacterial or archaeal pilus. We hypothesise that this feature may have important implications for the mechanism of twitching motility.

## Results

### CryoEM and helical reconstruction of Aap filaments

Filaments of *S. acidocaldarius* strains lacking archaella and Uvp (MW158) were sheared from cells at stationary phase and purified via CsCl gradient centrifugation. The resulting mix contained threads and Aap. The suspensions were plunge frozen on cryoEM grids, from which 6272 movies were recorded using a Titan Krios TEM. CryoSPARC ^45^ was used for the entire image processing workflow. The raw micrograph revealed highly flexible Aap filaments with curvatures of up to 90 degrees (Supplementary Figure 1). In contrast to the Aap, the archaella of the same organism show a slightly undulating superstructure and threads are mostly straight (Supplementary Figure 2).

Iterative 2D classification was used to separate the Aap from the thread filaments, the structure of which we solved previously ^3^. Using the Helix Refine function in CryoSPARC ^45^, we were able to reconstruct an initial unbiased 3.7 Å resolution map of the Aap without applying helical symmetry parameters (Supplementary Figure 3 b). Aided by the building of an initial atomic model, this map was sufficient to deduce the helical parameters, which we identified as a helical twist of -39° and a rise of 15.4 Å. Applying these parameters in subsequent refinements finally resulted in a map with 3.2 Å resolution (Figure 1, Supplementary Figure 3, 4 and 5). Our structure thus revises the previously published ∼ 9 Å resolution map of the *S. acidocaldarius* Aap ^29^, which was a result of the application of 136.9° twist and 5.7 Å rise.

**Figure 1.**
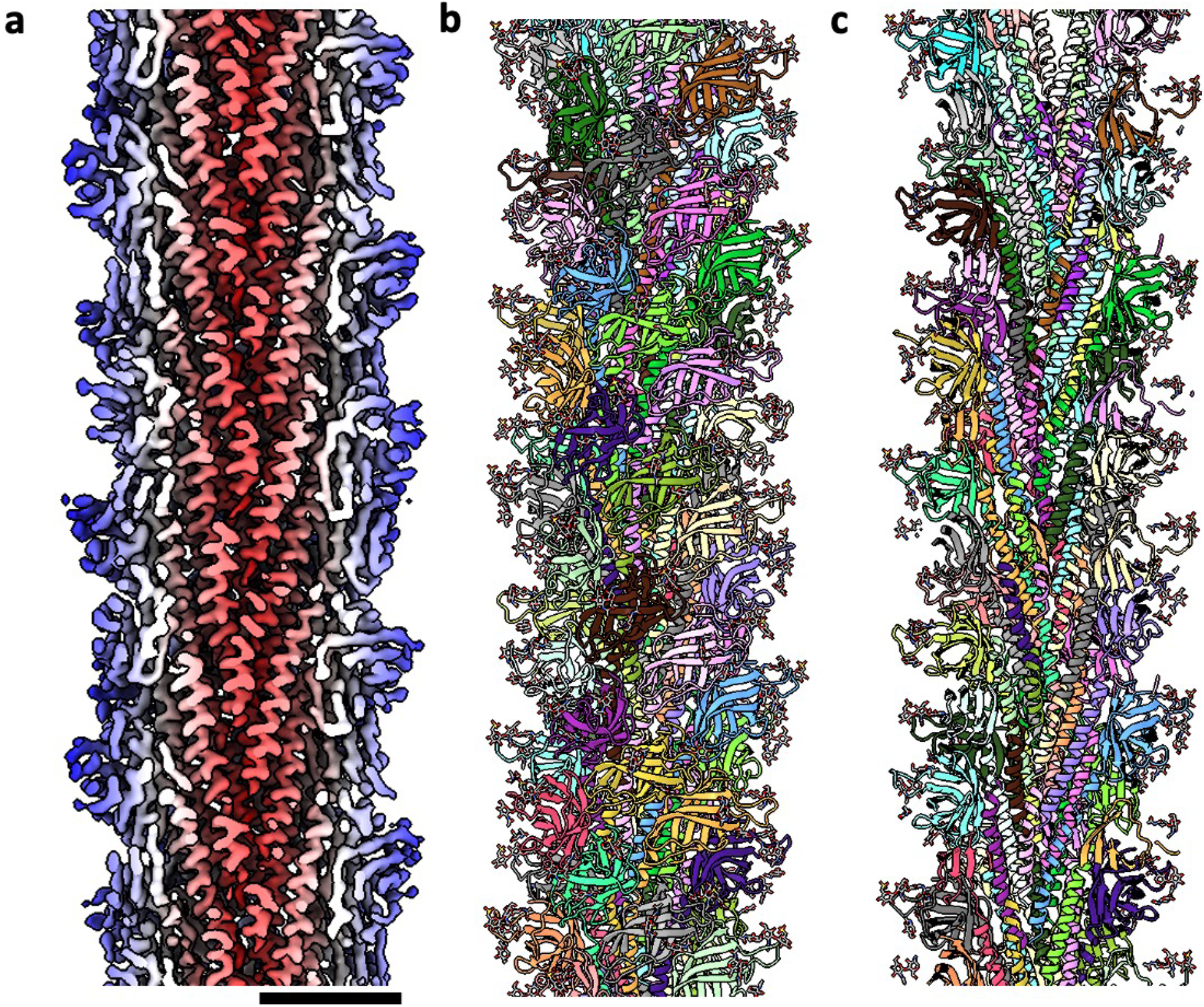
Helical reconstruction and atomic model of the *S. acidocaldarius* Aap **a,** Cross-section of the cryoEM map of the Aap, coloured by cylinder radius. The ⍺-helical core is highlighted in red, and the β-strand rich exterior in white and blue. **b,** atomic model of the Aap, with each subunit illustrated in a different color. Protein is shown in ribbon, and glycans are in stick representation. **c**, cross section of a, revealing the ⍺-helical core of the filament. Scale bar 60 Å

Moreover, the helical parameters identified by us differ from those published for homologous adhesive T4P of *S. islandicus* ^30^ and *S. solfataricus* ^31^, with previously published values around 105° twist and 5 Å rise. Indeed, applying similar values to our data resulted in a map with lower quality and resolution (Supplementary Figure 4). In addition, while the helical parameters of --39° twist and 15.4 Å rise enhanced the power spectrum of the refined map compared to the unbiased map, the FFT was scrambled when parameters close to the published 105° twist and 5 Å rise were applied (Supplementary Figure 3). While these erroneous helical parameters yielded a map where the ⍺-helices in the core of the filament were not particularly well defined, -39° twist and 15.4 Å rise greatly improved the secondary structure in the core, as well as the periphery of the filament (Supplementary Figure 3 and 4). Local resolution estimates showed that the core of the map has a resolution of 2.8 Å, whereas at the periphery of the filament, the resolution falls off to 3.8 Å (Supplementary Figure 5 b). The map clearly showed that the filament consists of multiple copies of tadpole-shaped monomers typical for T4P. In agreement with the resolution, large side chains could be clearly identified throughout.

### Composition and structure of the Aap

In *S. acidocaldarius*, the *aap* gene cluster encodes for two pilins, AapA and AapB (Supplementary 6 a). Previous experiments indicated that neither the deletion of *aapA* nor *aapB* led to loss of Aap filament formation ^29^. However, these findings were based on TEM of cells also producing archaella, and the low-resolution data precluded the visualisation of the differences between the two filaments. Re-evaluating this study, we found that the *ΔaapA* strain still assembles Aap (Supplementary Figure 7 a), whereas Aap were lost in the *ΔaapB* knockout (Supplementary Figure 7 b), as well as the double mutant *ΔaapAB* (Supplementary Figure 7 c).

We therefore asked if the filament would be composed of a mixed population of AapA and AapB, similar to the recently published structure of the archaellum of *Methanocaldococus villous* ^18^. By building the atomic model of the Aap pilus *ab initio* aided by large amino acid side chains, as well as glycosylation sites that differ in AapA and AapB (Supplementary Figure 8), we found that only the sequence of AapB could be reconciled with our map. This suggests that the filament is entirely comprised of AapB (Figure 1 b, c). We corroborated our findings by predicting the structure of AapB using Alphafold2. The prediction closely matched our *ab initio* model and thus confirmed AapB as the sole subunit composing the filament (Supplementary Figure 9). In line with these findings, mRNA sequencing data revealed that *aapA* is expressed in lower amounts than *aapB* ^46^ (Supplementary Figure 6 b). We thus suggest that this protein acts as a minor pilin, which may assemble into a cap or nucleating structure for the pilus. Alternatively, AapA could reside within the cellular membrane and function as a part of the Aap assembly machinery.

Each AapB subunit consists of an N-terminal α-helix tail, followed by a globular (head) domain. The head domain contains 8 β-strands that fold into two β-sheets (Supplementary Figure 10). T4P are N-terminally processed by a Class-III signal peptidase (PibD/ArlK in archaea) ^47^. In line with this, we noticed that 15 N-terminal amino acids were missing in the mature protein. Indeed, the FlaFind server ^48^, predicted a peptidase processing site between A15 and L16. Careful analysis of the filament’s structure revealed that the AapB monomers exist in three distinct conformations (henceforth designated as conformations A, B and C.) These differ in the angle between the head and tail domain with values of 106°, 128°, and 120° for conformations A, B and C, respectively (Figure 2 a, b; Movie1). The RMSD between conformations A and B is the highest with a value of 2.41 Å, whereas the RMSD between conformations S and B, and B and C are 1.94 Å and 1.60 Å, respectively (Supplementary Figure 11).

**Figure 2.**
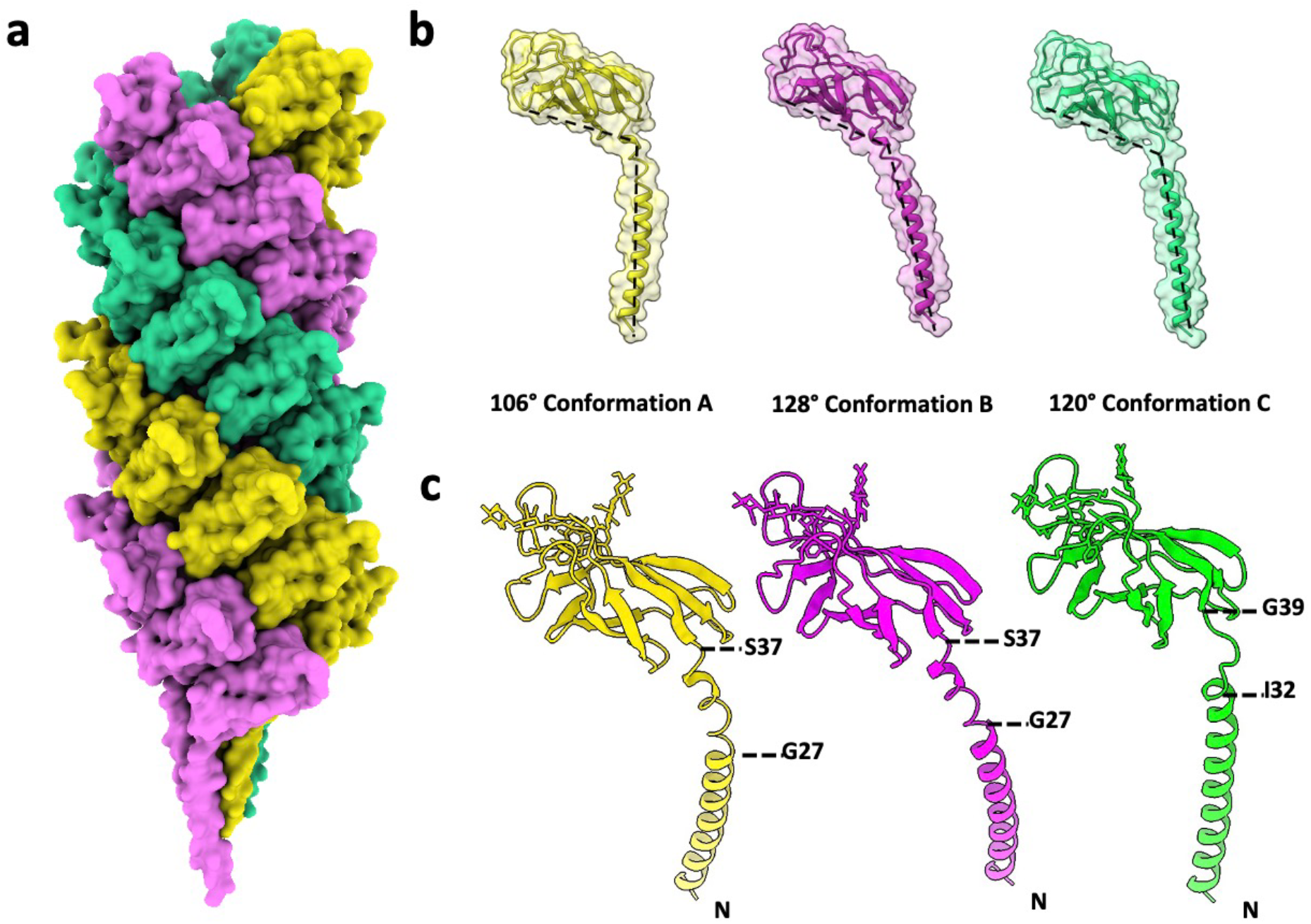
AapB adopts three conformations within the Aap **a,** structure of the *S. acidocaldarius* Aap in surface representation. The pilus is a 3-start helix and consists of three left-handed helical strands (magenta, yellow and green). Each of the three component helices is composed of multiple copies of AapB which adopts one of three conformations. **b,** AapB subunit in its three conformational states (named A, B and C). The colours of the three conformations indicate their location in the filament (a). The three conformations differ in the angle between the head and ⍺-helix tail (indicated by dashed lines), which is 106°, 128° and 120° for A, B and C, respectively. **c**, in conformations A and B, the ⍺- helix contains a distorted, partially melted stretch between G27 and S37. In conformation C, the ⍺-helix is fully melted between I32 to G39.

Closer inspection of the three structures revealed that the three distinct intra-subunit angles are due to differences in the melted region of the ⍺-helical tail. This region is a hallmark of all T4P and is characterised by a dissolution of the ⍺-helix into a loop of 3 to several amino acids in length ^49^. In conformation C of AapB, this region resembles the canonical melted loop of other T4Ps and lies between residues I32 and G39. However, in conformations A and B, the ⍺-helix does not appear to be fully melted. Starting at residue G27, the polypeptide transitions into a distorted ⍺-helix, which then links to the first β-strand (β1) following residue G37 (Fig. 2 c).

Interestingly, the conformation of AapB depends on its location in the filament. Each distinct conformation repeats along one of the three left-handed 3-start helices of the pilus, meaning that each of these 3-start helices harbours exclusively one of the three conformations (Figure 2 a). This leads to a structurally unique fibre that consists of three stacked helices with distinct conformations. Moreover, the ⍺-helical tails within the core are significantly curved, with the concave side of each ⍺-helix facing the filament’s periphery (Figure 3 a,d). This leads to an obvious screw-like distortion of the filament’s core. As Alphafold predictions of AapB suggest a straight ⍺-helix (Supplementary Figure 9), it is possible that the N-termini of AapB become bent upon integration into the pilus. Notably, a similarly distorted organisation is seen in the previously solved structure of the *S. islandicus* LAL14 pilus ^30^ (Figure 3b,e). In archaella, the archaellin subunits are slightly bent the other way, with the convex sides of the ⍺-helices facing towards the filament’s periphery (Figure 3 c, i) ^14, 16–18^. Despite this, there is no obvious screw-like distortion of the core of the assembled archaellum (Figure 3 f). In addition, while the heteropolymeric archaellum of *M. villosus* consists of two alternating subunits with a distinct sequence and glycosylation pattern (ArlB1 and ArlB2), the two subunits do not differ in the angle between head and tail (Figure 3 i). Thus, the distorted, tri-conformer organisation appears to be a unique feature of the *S. acidocaldarius* Aap.

**Figure 3.**
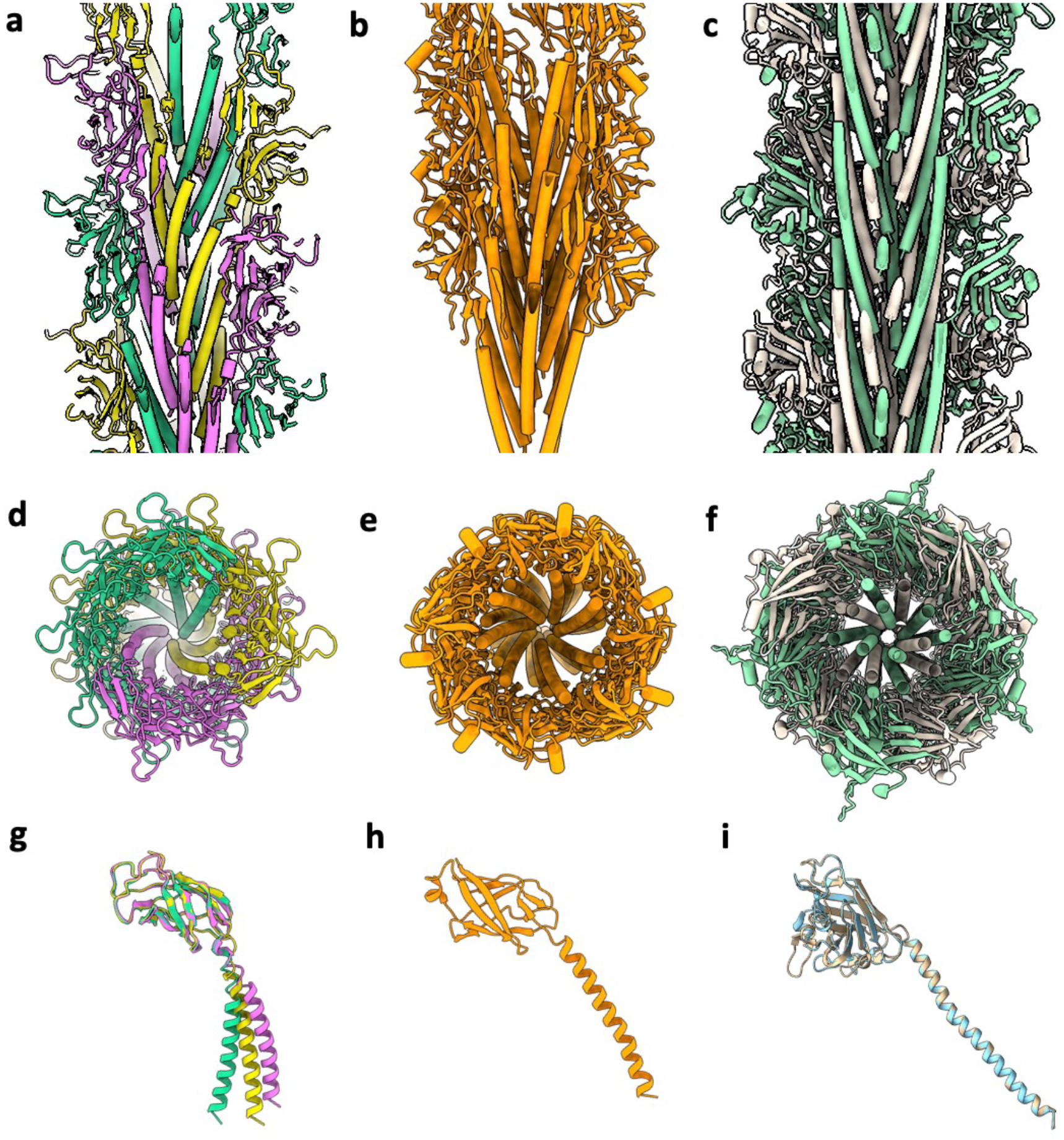
The *S. acidocaldarius* Aap in comparison with pili and archaella Longitudinal cross sections (**a-c**), top views (**d-f**) and subunits (**g-i**) of the *S. acidocaldarius* Aap (**a**,**d**), *S. islandicus* LAL14 filament (**b**, **e**; PDB-6NAV ^30^), and *M. villosus* archaellum (**c**, **f**; PDB-7OFQ ^18^). In the Aap (**a**, **d**), the central ⍺-helices are curved. Thus, the filament’s core adopts a screw-like architecture. A similar, albeit less well pronounced pilin organisation was seen in the *S. islandicus* pilus (**b**, **e**). In contrast, the archaellins in archaella appear rather straight and thus do not adopt a screw-like organisation. **g-i**, subunits of the *S. acidocaldarius* Aap (**g**), *S. islandicus* LAL14 filament (**h**) and *M. villosus* archaellum (**i**). Overlaying all three conformations of AapB highlights the differences in the three conformations. The *S. islandicus* LAL14 filament apparently only harbours one subunit conformation (**h**), however, the subunit is also slightly bent. **i**, while the archaellum of *M. villosus* is composed of two alternating subunits (ArlB1 and ArlB2), they only differ in the structure of the head domain, but not in the overall conformation of the subunits.

**Figure 4.**
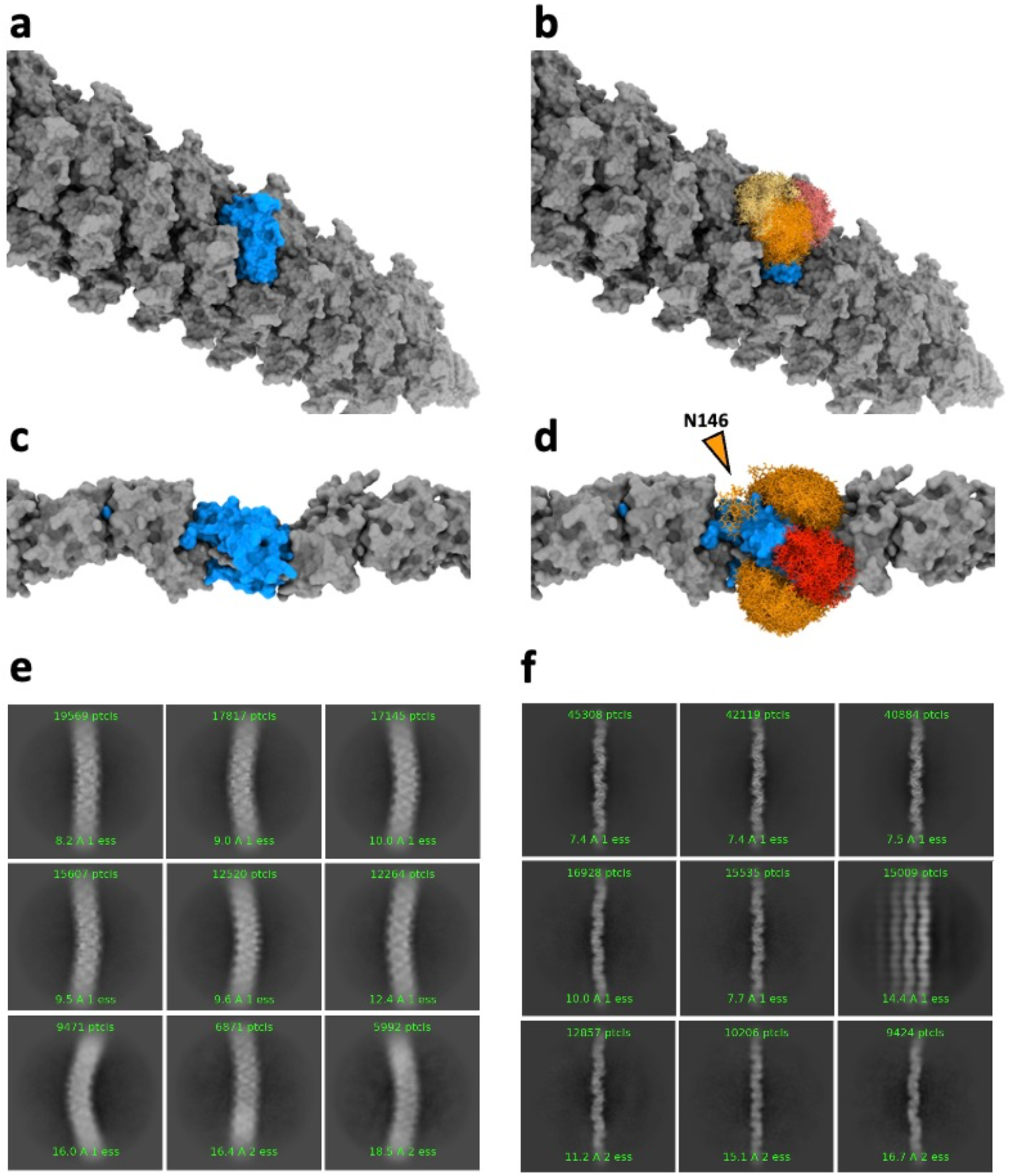
Glycosylation and flexibility of Aap compared to threads of *S. acidocaldarius* a, b. Glycoshield simulation ^55^, showing the large conformational space that the surface glycans (yellow, orange, red) occupy in a singular Aap subunit (a without and b with glycans for comparison). **c, d,** Glycoshield simulation for the four glycans in a singular *Saci* Thread subunit (c without and d with glycans). In contrast to the Aap, one of the thread glycans (N146, orange arrowhead) occupies a limited conformational space, as it is wedged between the interface of two subunits. **e, f,** 2D classes of Aap (**e**) and threads (**f**) show differences in stiffness between both filaments.

### The Aap is posttranslationally modified with highly flexible N-glycans

As we were building the atomic model of the Aap, we identified three glycans per AapB subunit. In the cryoEM map, these became apparent as dead-end protrusions which which had characteristic features of archaeal glycans ^3, 16, 18^. The glycan densities originated from asparagine residues, (N48, N60, and N88), which were all part of a consensus N-glycosylation sequon (NXS/T). In *S. acidocaldarius,* N-linked glycosylation utilises tri-branched hexasaccharides, featuring the rare 6-sulfoquinovose sugar. We previously solved the structure of this glycan by combining our structure of the *S. acidocaldarius* thread filament ^3^ with previously published mass spectrometry data ^50–52^, and modelled this structure into the glycan density of the Aap (Supplementary Figure 12 b-e). Notably, we found a fourth Asparagine with the N-glycosylation consensus sequon located at N114, however, no glycan density was present at this residue. This differs from our structures of the *S. acidocaldarius*, thread and *M. villosus* archaellum, where every surface-exposed N-glycosylation sequon is glycosylated ^3, 18^. The absence of the glycan at position N114 may be explained by potential inaccessibility of the site in the preprotein, or previous findings, suggesting that AglB, the enzyme that glycosylates proteins in *Sulfolobus*, is promiscuous ^53, 54^. When comparing the glycan densities of the Aap pilus with those of the thread filament, it becomes clear, how much better defined the glycans are in the threads (Supplementary Figure 12 b-j) ^3^, even though both filaments were resolved at similar global resolution and using the same sample. While all five sugar units of the hexasaccharide were resolved in the threads, the map of the Aap pilus contained only contained density for the first sugar residue (Figure 12 a-e). Densities for the two N-acetylglucosamine (GlcNAc) residues, the connecting mannose (Man), glucose and 6- sulfoquinovose molecules were not resolved, reminiscent of previous maps of T4P, of *P. arsenaticum, S. solfataricus* ^31^, and *S. islandicus* ^30^. Asking whether this difference is due to various degrees in glycan flexibility, we performed molecular dynamics simulations (Figure 4). This analysis revealed that the glycans in the thread indeed adopt fewer conformations than in the Aap pilus. This is particularly true for the tread glycan bound to Asn146, which is wedged inside a cleft between two adjacent thread subunits. Because of this reduced mobility of thread glycans, their structure can be reconstructed in helical processing, while the highly flexible Aap glycans largely average out.

### Filament flexibility

It has previously been suggested that the flexibility of glycans correlates with the flexibility of proteins ^55^, suggesting that Aap should be more flexible than threads. Indeed, our cryoEM micrographs (Supplementary Figures 1 and 2), as well as 2D classes (Figure 4 e,f) indicate that threads are usually straight, while Aap are able to bend considerably. To understand the structural basis of the apparent differences in filament stiffness, we performed 3D Variability Analysis (3DV) in CryoSPARC and built atomic models for the generated output.

While this 3DV approach could only probe a limited conformational range, the analysis confirmed our 2D classification data. Aap appear more flexible compared to the threads of the same organism (Supplementary movie 2). This is likely functionally important, as threads are mainly thought to be important for cell-cell adhesion and biofilm formation ^3^, while Aap are involved in twitching motility ^32^.

These differences in flexibility can be explained with the distinct intermolecular interactions that hold Aap and threads together. In the Aap, the main intermolecular contacts are found in form of hydrogen bonds between the c-terminal and hydrophobic interactions between the N- terminal tails. As previously shown for the archaellum, the Aap heads are linked to the tails by a conserved hinge region and are thus free to move with respect to the tails ^18^. In the Aap, this hinge region is pronounced, and depending on the conformational state of AapB, encompasses up to 10 residues. It is intriguing to speculate, that the distinct melting states of the hinge regions may lead to different stiffnesses of the 3 component helices. In addition, the hydrophobic interactions in the core of T4P and archaella act as molecular grease that enables the tails of the subunits to slide past each other ^18^, and this can be seen in the 3DV analysis of the Aap (Supplementary movie 2). The threads, on the other hand, do not have a hydrophobic core that could provide the same degree of flexibility. Instead, the subunits are interlinked by donor strand complementation (DSC), where the N-terminal tail of one subunit integrates into and completes the β-sheet of the next subunit (N+1) along the chain ^3^. β-sheets are rich in hydrogen bonds and thus do not allow for significant flexibility. Furthermore, there are no pronounced intramolecular hinges in the thread and an isopeptide bond is formed between subunit N and N+2 ^3^, covalently fixing the N-termini of the protomers within head subunits. Furthermore, the glycan attached to N146 of the thread, which is wedged into a cleft between two adjacent subunits (Figure 4 f, Supplementary Figure 12 j), likely restricts the movement of the subunits with respect to each other. No sterically wedged glycans occur in the Aap. Instead, the glycans are free to explore a large range of conformations, thus allowing for a highly flexible Aap filament.

### Aap are conserved among some Crenarchaea

To investigate the conservation of Aap, we used SyntTax to search for AapB homologues in related archaeal Thermoprotei species and found six highly conserved proteins (Supplementary Figure 13). Henche et al. ^29^ previously suggested that the two of these species (*S. islandicus* and *S. solfataricus*) would not assemble Aap, as their AapB homologs are not located near the machinery genes. However, in recent years, the structures for both these Aap filaments were solved ^30, 31^. The helical parameters for both pili were determined at 5 Å rise and ∼105° twist, in contrast to our structure with 15 Å rise and -39.9° twist. Despite this, the all identified homologs are highly conserved on the amino acid level, and so we asked if structural differences were evident in comparison with our tri-conformer Aap filament from *S. acidocaldarius*.

Multisequence alignments revealed almost identical N-terminal ⍺-helices in all identified homologs, as well as several key glycine residues throughout the entire sequence. In all cases, Alphafold2 suggested almost identical structures when compared to *S. acidocaldarius* AapB (Supplementary Figure 13 a,b). Apart from the N-terminal tails, regions of high similarity included the core-facing β-strands of the head region, with greater variability in the solvent-exposed strands and the loops connecting them. The nearly interchangeable alpha helices indicate that these homologues may in fact all follow the helical parameters identified for the *S. acidocaldarius* Aap. Thus, the homologs may in fact also form pili that adopt similar tri- conformer structures, as it is the pilin tails that dictate this feature. In contrast, nuances such as the glycosylation sites, which are highly variable (Supplementary Figure 13 d), suggest that the position (and likely sequence) of the glycans play a key role in adapting each species to its environment or function. The widespread nature of the Aap raises the question if twitching motility is equally widespread in archaea.

### A model of the Aap Machinery

*AapB* is encoded in a gene cluster, together with putative machinery components *aapX, aapE and aapF* (Supplementary figure 6 a). We used Alphafold2 to predict the structures of these proteins and, together with our solved structure of the Aap, integrated them into the current working model of the Aap machinery (Figure 5 a). AapF has previously been shown to be partially homologous to the membrane integral T4P platform protein PilC from *T. thermophilus* ^56^, (Supplementary Figure 14 a), as well as the archaellum assembly platform ArlJ from *S. acidocaldarius* ^57^, (Supplementary figure 14 b). ArlJ is predicted to be a dimeric protein ^57^, and so we modelled AapF in the same manner. Deep TMHMM predicted that AapF is a 9- transmembrane helix protein, with a bulky N-terminal domain in the cytoplasm and short periplasmic loops (Supplementary figure 15). Comparing AapF with ArlJ showed the greatest variability in the N-terminal region, suggesting a potential role in determining the proteins’ substrate selectivity (pilin for ArlF vs. archaellin for ArlJ) (Supplementary figure 14 e,f).

**Figure 5.**
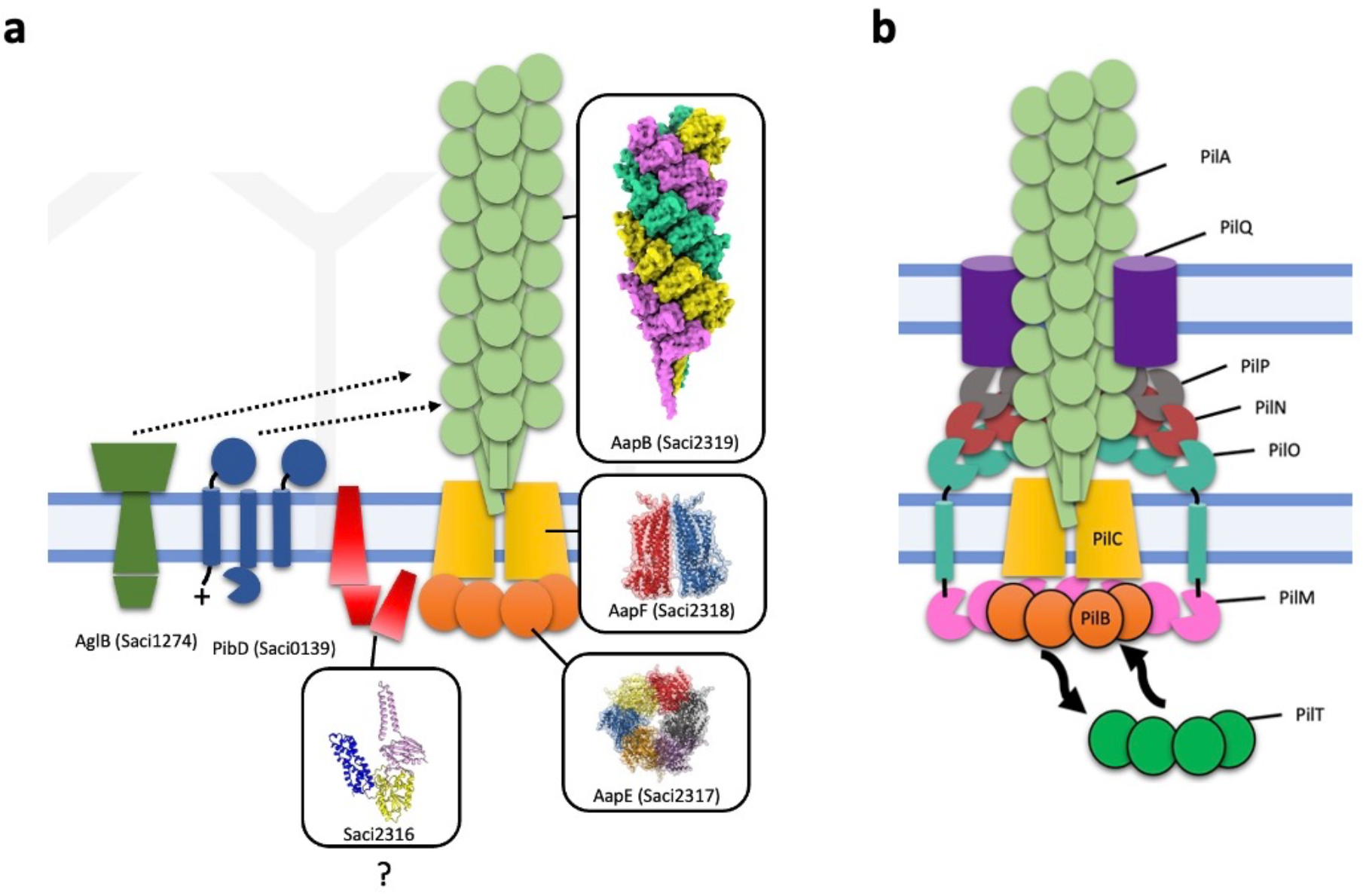
Model comparing the archaeal and bacterial twitching apparatus **a,** model of the Aap machinery from *S. acidocaldarius*. The pilus is composed of AapB and assembled by the putatively dimeric, and membrane-integral platform protein AapF. The latter binds to the cytoplasmic ATPase AapE. AapX is also likely part of the machinery, but its function is unknown. The signal peptidase PibD is responsible for the cleavage of the AapB pre-pilins, which primes them for pilus integration. AglB is the enzyme that glycosylates cell- external proteins, including AapB. AapB pilins assemble into the tri-conformer pilus. If the pilins adopt their three conformations prior to or during pilus assembly is unknown. **b,** model of the bacterial T4P machinery from *Neisseria gonorrhoeae*, ^87^. The PilA filament is assembled by the membrane-integral platform protein PilC and transverses through a PilQ pore, which is integrated within the outer membrane and peptidoglycan layer. Within the cell wall, rings of accessory proteins (PilO, N and P) surround the filament. PilO extends down through the cell membrane to bind a ring of PilM, which encompasses the pilus assembly ATPase PilB. The latter is substituted for the disassembly ATPase PilT to rapidly retract the pilus and thus drive twitching motility. So far, only single conformer pili are known in bacteria.

AapE shows sequence similarity with the archaellum assembly ATPase ArlI (Supplementary Figure 16 a) ^58^ . This similarity is reflected by the Alphafold2 prediction for AapE, which is consistent with the structure of the AAA ATPase (Supplementary Figure 16 b). Indeed, it has previously been shown that the homolog from *S. solfataricus* functions as an ATPase ^58^. As is common for AAA ATPases, ArlI is a hexameric protein ^59^, and so we modelled AapE as a hexamer (Supplementary Figure 16 c,d). Homologues of AapE and AapF were found in all six archaeal strains that also encode for an Aap-like pilin. Both genes always appear next to each other in the genome, although not necessarily near to the subunit homologue of AapB ^29^.

Alphafold2 predictions and PDBe-fold analysis of AapX indicated a membrane-anchored protein (Supplementary figure 17). Homologs of the *aapX* gene are also found in all other organisms that contain an *aapB* homolog and are located ∼≤10 genes upstream from the putative ATPase and platform protein counterparts ^29^. Interestingly, AapX has a transmembrane helix, which is absent in the other species investigated. Instead, these homologues have an FAD binding domain (Supplementary Figure 17 d,e), which is not present in *S. acidocaldarius.* As with *aapE* and *aapF* mutants, knockouts of *aapX* do not express Aap filaments ^29^. Thus, while the function of AapX is unknown, it is necessary for the Aap assembly and likely part of the Aap machinery.

## Discussion

Here we present the structure of the Aap pilus from *S. acidocaldarius* at 3.2 Å resolution. We find that the subunits of each 3-start helix exist in a distinct conformation. Conformational variability on the subunit level has recently been observed in flagella and archaella ^20^, as well as the bacterial flagellar hook of *Salmonella enterica* ^60^. The FlgE hook of *S. enterica* consists of 11 subunits per turn, with each monomer inheriting a subtly different conformation that leads to the observed bending of the hook ^61^. Similarly, it has been suggested that in archaellar and flagellar filaments from various species, conformational differences in the subunits generate the supercoiling necessary for swimming propulsion ^18, 62^. Whilst these observations explain how filaments adopt minimum energy states in clockwise vs anticlockwise rotation, the conformations found here in the Aap pilus are entirely different and unique. In the Aap, we observe a 3-start helix configuration, where each of the 3 component helices adopts an independent structure that is consistent through the entire length of filament. The three conformations are largely established through different extents of melting of the hinge region of the AapB subunits.

In bacteria, pilus retraction is thought to be triggered as the pilus touches a surface. Presumably, a retrograde signal then ripples from the distal adhesion point to the pilus’ assembly machinery in the cell envelope ^63, 64^. This activates pilus retraction, whereby the filament disassembles into its constituent subunits, which concomitantly transfer from the disassembling filament into the cell membrane ^63–65^. For efficient twitching, inter-subunit interactions within pili must be sufficiently strong to enable the assembly of a pilus, but also sufficiently weak, to allow swift disassembly during retraction. Thus, pili that are involved in twitching must exist in a metastable state.

Indeed, point mutations that weaken the subunit interactions between the major pilins of *Vibrio cholerae* decrease the stability of the filament. This allows pili disassemble easier and thus to retract quicker than the wild type, even without the aid of any ATPase activity ^66^. The unusual triple conformer helix of *S. acidocaldarius* Aap may have evolved to aid pilus retraction.

Notably, the three distinct conformations of the pilus evoke a twisted filament core, in which the ⍺-helices are unusually bent. Moreover, the three pilins in the Aap exist in differently bent conformational states, as opposed to a straight conformation suggested by Alphafold2 (Supplementary Figure 9), or as seen in previously solved structures of archaella, where the core organisation is virtually straight. As the lowest energy of an ⍺-helix is when it lies in a straight conformation, the monomers within the Aap likely exist in an elevated energy state, comparable to a loaded spring. It is conceivable that this energy could be spontaneously released (for example as the pilus touches a surface), thus eliciting the elusive retrograde signal through the pilus, or its collapse into the cell membrane. The latter could exert a force capable of pulling the cell forward. In accordance with this hypothesis, in both *P. aeruginosa* and *N. gonorrhoeae* ^67^ AFM-induced tension resulted in conformational changes in the filaments. The distinctive structure of the Aap also necessitates heterogeneous interfaces between the three different pilin conformations. Applying the helical parameters that were published for the pilus from *S. solfataricus* and *S. islandicus* ^30, 31^, results in a pseudo- homogeneous structure with of a straight core (Supplementary Figure 4 a). Such a hypothetical organisation would lead to a closer packing between the central ⍺-helices of AapB than seen in the heterogenous Aap pilus, and thus would likely render pilus disassembly less efficient due to stronger interactions.

Delving into the evolution of *S.acidocaldarius* Aap from the last universal common ancestor, the closest related group of bacteria to archaeal species are those that produce tad pili ^11^. The full list of 14 genes needed for a fully functional tad pilus far exceeds those needed for the Aap ^68^, and so the support systems surrounding the filaments differ greatly, likely due to the difference in cell walls (Figure 5 b). Notably, the Tad pili, along with gram-positive competence pili, Type II secretion filaments and some Type IVb filaments all possess only one ATPase, yet are all capable of filament retraction ^69^. The Tad pilus ATPase in *C. crescentus,* CpaF, has been shown to hydrolyse the extension and retraction of tad pili – a mechanism that was also found in other species ^69^. This finding disproved the previously hypothesised retraction mechanism centred around the minor pilins encoded in the pilus operon, showing that the minor pilins may be dispensable for pilin formation, within retraction-deficient backgrounds ^44^. Likewise, only one type of ATPase can be found for the Aap system of *S. acidocaldarius*, as well as related archaeal species, and only a single ATPase is currently known to drive the assembly and rotation of the archaellum, both in clockwise and anticlockwise rotation. It therefore appears plausible that Aap assembly and retraction could also occur through the bifunctional ATPase, in *S. acidocaldarius* and related species. Furthermore, based on the Aap gene cluster, the assembly machinery appears to be far simpler than that of its bacterial counterpart (Figure 5). Whereas the bacterial T4P complex encodes for the alignment complex PilMNOP (Figure 5 b) and a secretin (PilQ) in diderm bacteria, the crenarchaeal Aap pilus lacks these components (Fig, 5 a). It is conceivable that this alignment complex is obsolete, as *S. acidocaldarius* does not have a second membrane. Instead, the Aap must traverse the S-layer - a porous, proteinaceous cage that surrounds many archaea. Whether a so-far elusive protein complex is required to guide the Aap through the S-layer or whether the S-layer itself acts as a guiding scaffold remains to be elucidated.

## Materials and Methods

### Cell growth and Aap isolation

Pilus isolation was performed as described in Gaines et al., 2022 ^3^. Briefly, MW158 was inoculated from cryo stock into 6 x 5 ml basal Brock at pH3, supplemented with 0.1 % NZ-amine, 0.2 % dextrin and 10 µg/ml Uracil. Cell growth was carried out for 48 hours at 75 °C accompanied by light agitation. 5 mL of the resulting preculture was inoculated per litre of main culture and cells were grown to OD600nm = 0.5-0.8. Centrifugation at 5000 x *g* was then carried out for 25 mins at 4 °C to harvest the cells. Cell pellets from this 2 l main culture were then resuspended in 20 ml Basal Brock (pH 3) without FeCl3. Aap filaments were sheared as described in Henche et al 2012 ^29^, or via a peristaltic pump (Gilson Minipuls), connected to a syringe needle with 1.10 mm in diameter and 40-50 mm in length (Braun GmbH). Homogenisation at 25 rpm was imposed for 1 h before the syringe needle was switched to narrower ones with 0.45 mm and 0.10 mm diameter for shearing at 25 rpm for 1 h. Sheared samples were then centrifuged at 12 000 x *g* for 25 min at 4 °C followed by a subsequent ultracentrifugation at 200 000 x *g* for 90 min at 4 °C. This pellet was then resuspended in 500 µl Basal Brock without FeCl3 and layered on 4.5 ml CsCl (0.5 g/mL). Density gradient centrifugation was done at 250 000 x g for 16 h at 4 °C and the resulting white band in the upper third of the tube was collected. This band was diluted with 5 ml of Basal Brock without FeCl3 and pelleted at 250 000 x *g* for 1 h at 4 °C. Resuspension into 150 µl of citrate buffer (25 mM sodium citrate/citric acid, 150 mM NaCl, pH 3) was done before storage at 4 °C.

### Deletion of genes in S. *acidocaldarius*

Used Strains:

| Strain | Genotype | Source/Reference |
| --- | --- | --- |
| **MW158** | Deletion of *upsE* and *flaJ* in MW001 | Henche et al. 2012<sup>29</sup> |
| **MW153** | Deletion of *aapA* in MW001 | Henche et al. 2012<sup>29</sup> |
| **MW154** | Deletion of *aapB* in MW001 | Henche et al. 2012<sup>29</sup> |
| **MW161** | Deletion of *aapA* and *aapB* in MW001 | Henche et al. 2012<sup>29</sup> |
| **MW114** | Deletion of *pibD* in MW001 | Henche et al, 2014<sup>8</sup><br>Gaines et al 2022<sup>3</sup> |
| **MW2108** | Deletion of *saci_0405-0406* in MW114 | Gaines et al 2022<sup>3</sup> |

### Negative stain transmission electron microscope

Freshly glow-discharged 300 mesh carbon coated copper grids (Plano GmbH, Wetzlar Germany) were used to apply 5 µl of *S. acidocaldarius* cells, which were then incubated for 30 s. Excess liquid was blotted away and cells were re-applied on the grid. This was repeated three times. Grids were stained with 2% Uranyl acetate. A Hitachi HT8600 operated at 100kV equipped with an EMSIS XAROSA CMOS camera was used for imaging.

### Cryo-EM sample preparation and data collection

3 μl of a heterogenous population of Aap and threads was pipetted onto glow-discharged 300 mesh copper R2/2 Quantifoil grids. Blotting was performed using 597 Whatman filter papers for 5 s combined with a blot force of -1 in an environment of 95 % relative humidity and 21 °C. A Mark IV Vitrobot (FEI) was used to plunge-freeze the sample into liquid ethane. Grid screening was performed in a 120 kV FEI Technai Spirit EM combined with a Gatan OneView CMOS detector (Gatan, Pleasanton, USA).

For high resolution image data collection, a FEI Titan Krios electron microscope (Thermo Fisher Scientific, Eindhoven, The Netherlands), operating in nanoprobe mode using parallel illumination and coma-free alignment at a voltage of 300 kV, combined with a Gatan K2 Summit electron detector (Gatan, Pleasanton, USA) was used. The detector was operated in counting mode, at a calibrated magnification of x 134,048 (relating to a pixel size value of 1.047 Å). EPU software (Thermo Fisher Scientific) was used to control the camera. Movies were recorded at a dosage of 0.77 e/Å2 s−1 at 40 frames s−1, 10 s exposure, with an accumulated total dose of 42.33 e/Å2 and a set defocus range of -2.0 to -3.5 µm, using 0.3 µm steps. Cryo-EM statistics are shown in Table 1.

### Cryo-EM image processing

The Full-frame motion correction package from cryoSPARC ^45^, was used to align the frames of 6272 movies. Defocus variation was then estimated using the patch CTF estimation program. The e2helixboxer program from EMAN2 ^70^, was used to manually pick Aap filaments to generate basic 2D classes in Relion. These classes were then imported to cryoSPARC, so the Filament tracer program could pick out fragments comprising of Aap filaments. Originally only the best 2D classes were picked, the quality of the unbiased 3D helix refine job remained poor, likely due to a limited range of projections. To overcome this, a larger number of 2D classes were picked, aiming to include more views compromising for resolution. Using these 2D classes a total of 947 729 particles was subjected to non-biased 3D refinement, resulting in map with 3.74 Å resolution. Using this map, the helical parameters were determined at 15.4 Å rise and -39.9° twist. Local motion correction and CTF refinement as well as global CTF refinement jobs were carried out before a final helix refine job, which reached a global resolution of 3.22 Å. This resolution was estimated using Fourier shell correlation between two independently refined half sets, using the Gold Standard correlation value of 0.143. The map was denoised and postprocessed using DeepEMhancer ^72^. Local resolution was calculated within cryoSPARC ^45^, and maps were visualised in ChimeraX ^71^.

### Model building and validation

Initial manual model building in Coot enabled the unambiguous identification of AapB as Aap subunit. This was possible due to the glycosylation pattern of AapB, which differs significantly from AapA. This assignment was corroborated by comparing it to the predicted structure determined using AlphaFold2 ^73^. MOLREP ^74^, was then used for phased molecular replacement, to position the remaining monomers into the density. Changes in the rebuilt model were propagated using CCP4 ^75^, so that the copied monomers fit into the other densities. Coot was used to model the glycan structures ^76^, with unusual sugars dictionary being prepared using the JLIGAND ^77^. Refinement of the final structure was done using REFMAC5 via the ccpem interface ^78, 79^.

### Sequence analysis and structural prediction

The AapB sequence was loaded into SyntTax ^81^, in order to search for homologous sequences amongst other Thermoprotei species. The top six differing results were then compared against each other using Clustal Omega ^82^. To visualise the predicted gene cluster surrounding AapB, the KEGG genome database was used ^80^, followed by Synttax to search for homologous proteins in differing archaeal species ^81^. Protein structure prediction was performed with AlphaFold2, using the online ColabFold tool ^83^.

### 3D variability analysis and molecular flexibility

The flexibility of the Aap and threads was analysed using the molecular 3D variability tools in cryoSPARC ^45^. For both filaments, 20 frame flexibility modes were calculated and visualised in USFC Chimera ^84^. The mode showing the most significant flexibility was used to build atomic models. In brief, the refined models were positioned into fame 0 of the map series using MOLREP and the structures were then refined against the maps from the 1^st^, 5^th^, 10^th^, and 20^th^ frame of each map series in real space using Phenix ^85^. The subsequently generated maps and models were then used to create a morph in ChimeraX ^71^, showing the flexibility of the filaments.

### Data availability

The cryoEM map generated in this study has been deposited in the EM DataResource under accession code EMD-18119. The corresponding atomic coordinates have been deposited in the protein Data Bank database under accession code PDB-8Q30. The *S. acidocaldarius* (DSM639) genome can be accessed via the KEGG accession code T00251 (https://www.genome.jp/entry/gn:T00251) or the NCBI gene bank code CP000077 112. The transcriptomics data analysed in this study can be accessed in the Pan Genomic Database for Genomic Elements Toxic To Bacteria.

## Acknowledgements

We would like to thank Diamond Light Source for providing access to the cryoEM facilities at the UK national electron bio-imaging centre (eBIC), funded by the Wellcome Trust, MRC and BBSRC. Filament preparations were initially screened Faculty of Biology at the EM facility of the University of Freiburg, who we would like to thank for providing access to their microscopes. The TEM (Hitachi HT7800) was funded by the DFG grant (project number 426849454) and is operated by the University of Freiburg, Faculty of Biology, as a partner unit within the Microscopy and Image Analysis Platform (MIAP) and the Life Imaging Center (LIC), Freiburg. BD, MG, MM and RUH were supported by an ERC Starting under the European Union’s Horizon 2020 research and innovation program (grant agreement No 803894), awarded to BD. MM was also funded by a BBSRC New Investigator Research Grant (BB/R008639/1) to VG. SS and SVA were supported by the Collaborative Research Centre SFB1381 funded by the Deutsche Forschungsgemeinschaft (DFG, German Research Foundation)—Project-ID 403222702—SFB 1381. SS and SVA were also funded by the Deutsche Forschungsgemeinschaft (DFG, German Research Foundation) under Germany’s Excellence Strategy (CIBSS – EXC-2189 – Project ID 390939984). CH was supported by the Agence Nationale de la Recherche (grants #ANR-16-CE16-0009-01 and #ANR-21-CE16- 0021-01).

