## Supplementary figures for "CryoEM reveals the structure of an archaeal pilus involved in twitching motility"

#### Supplementary Figure 1

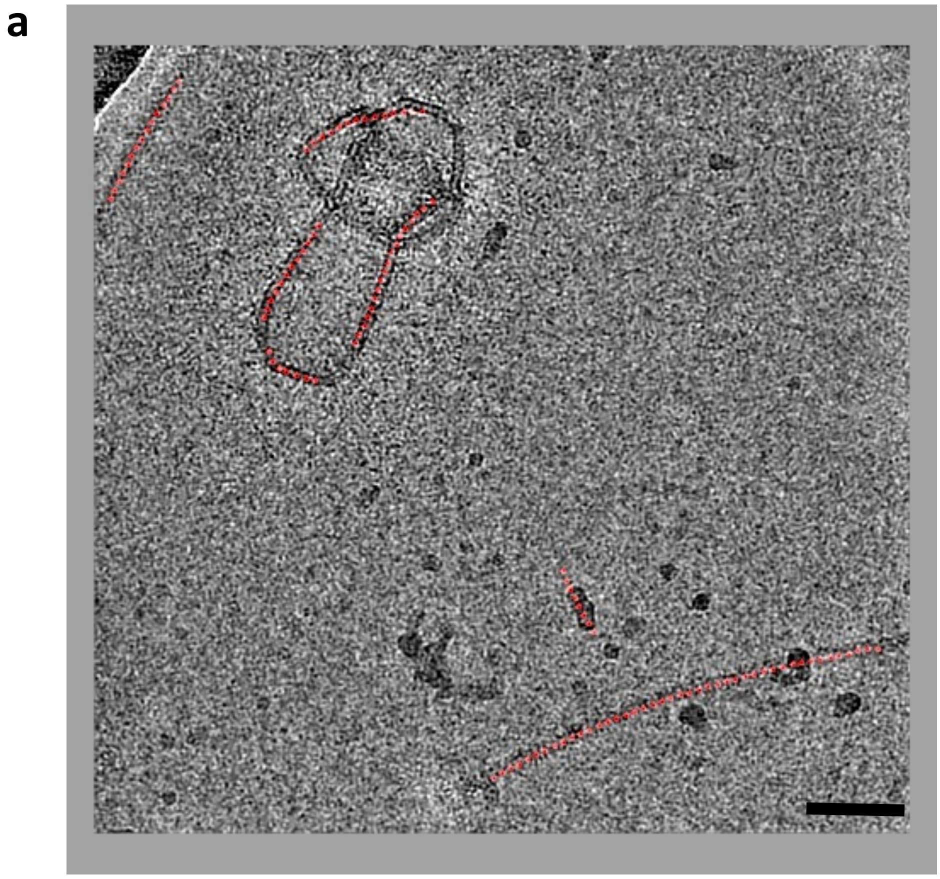

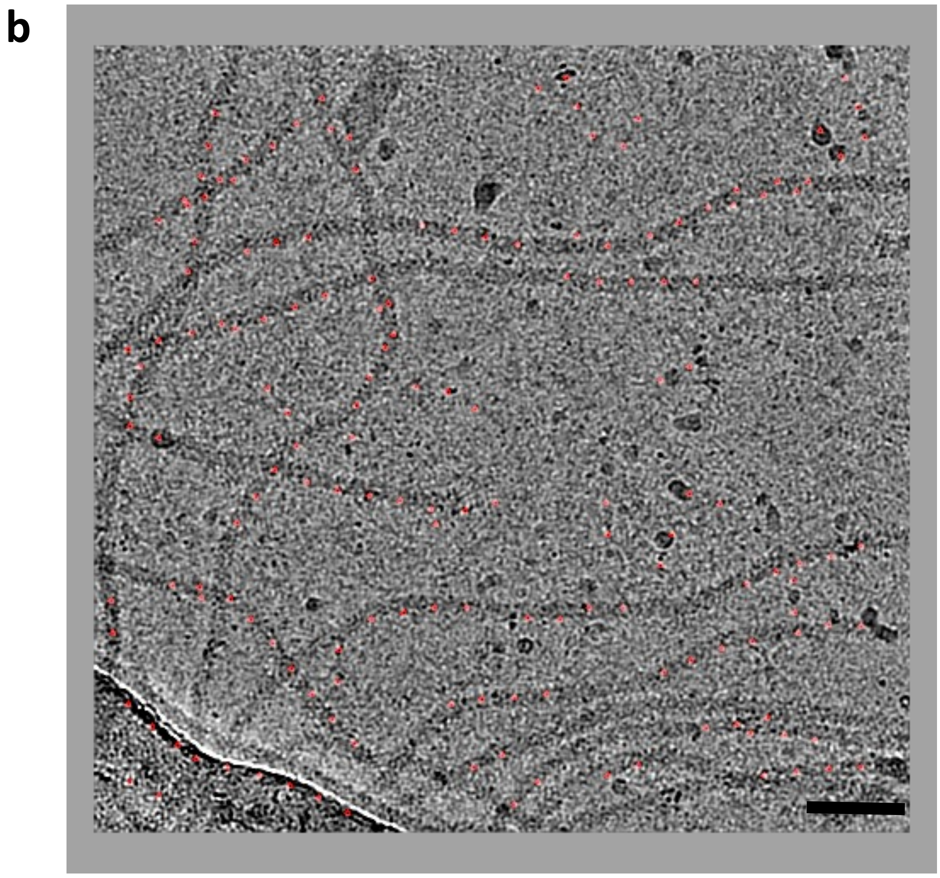

**Supplementary Figure 1 – CryoEM of isolated Aap**

**a, b,** raw micrographs of isolated and vitrified Aap indicate that Aap can adopt high degrees of curvature.

Scale bar 500nm

#### Supplementary Figure 2

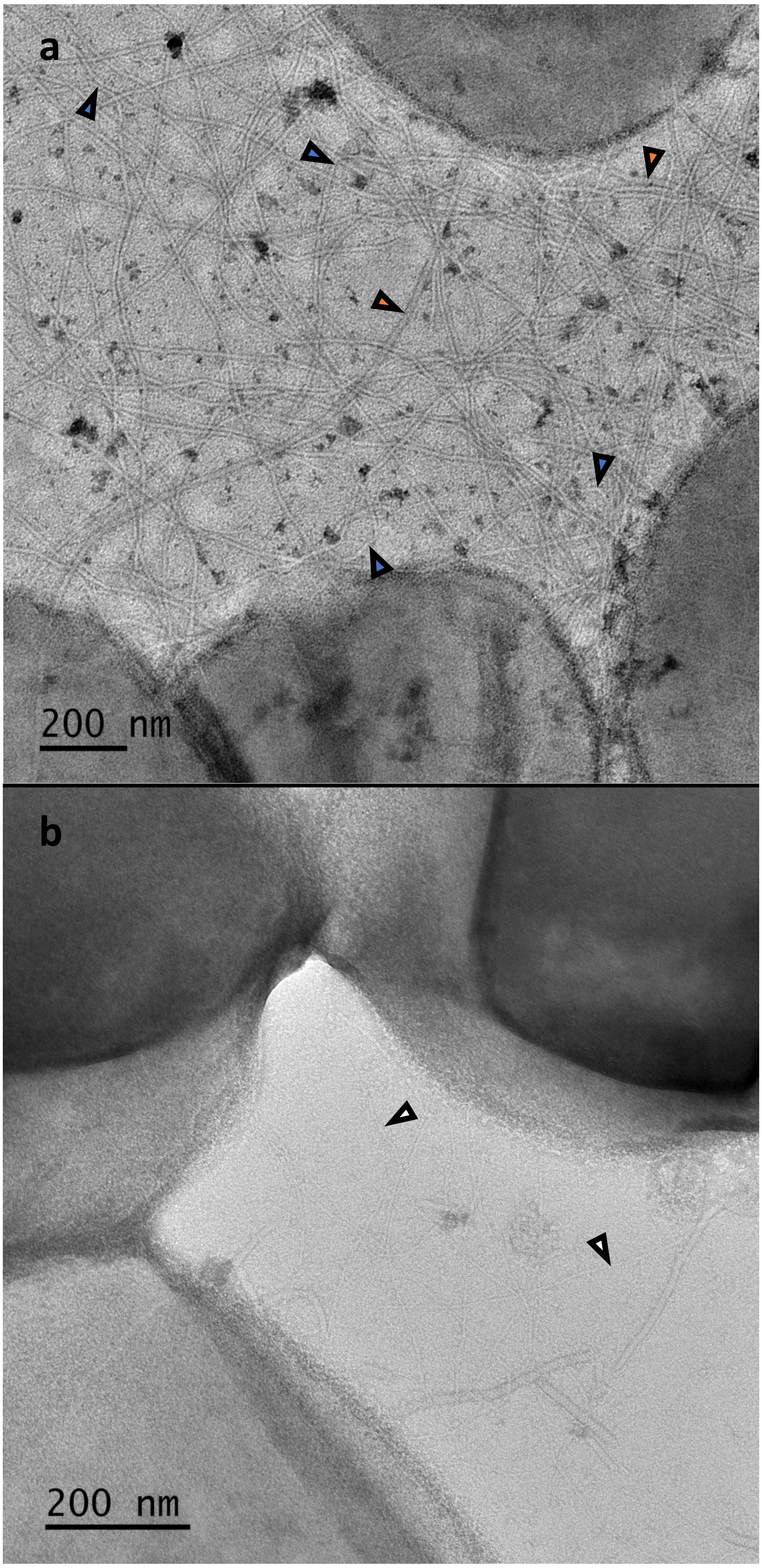

**Supplementary Figure 2 – Negative stain EM of *wt S. acidocaldarius***

**a, b,** in electron micrographs of negatively stained cells, Aap (blue arrowheads; a and b), archaella (orange arrowheads; a and b) and threads (white arrowheadss; b) can be distinguished by diameter and apparent stiffness. Aap have a diameter of ~8 nm[2] and are highly curved. Archaella are ~12 nm[1] wide and undulate slightly. Threads measure ~4 nm[3] in diameter and appear relatively straight.

### Supplementary Figure 3

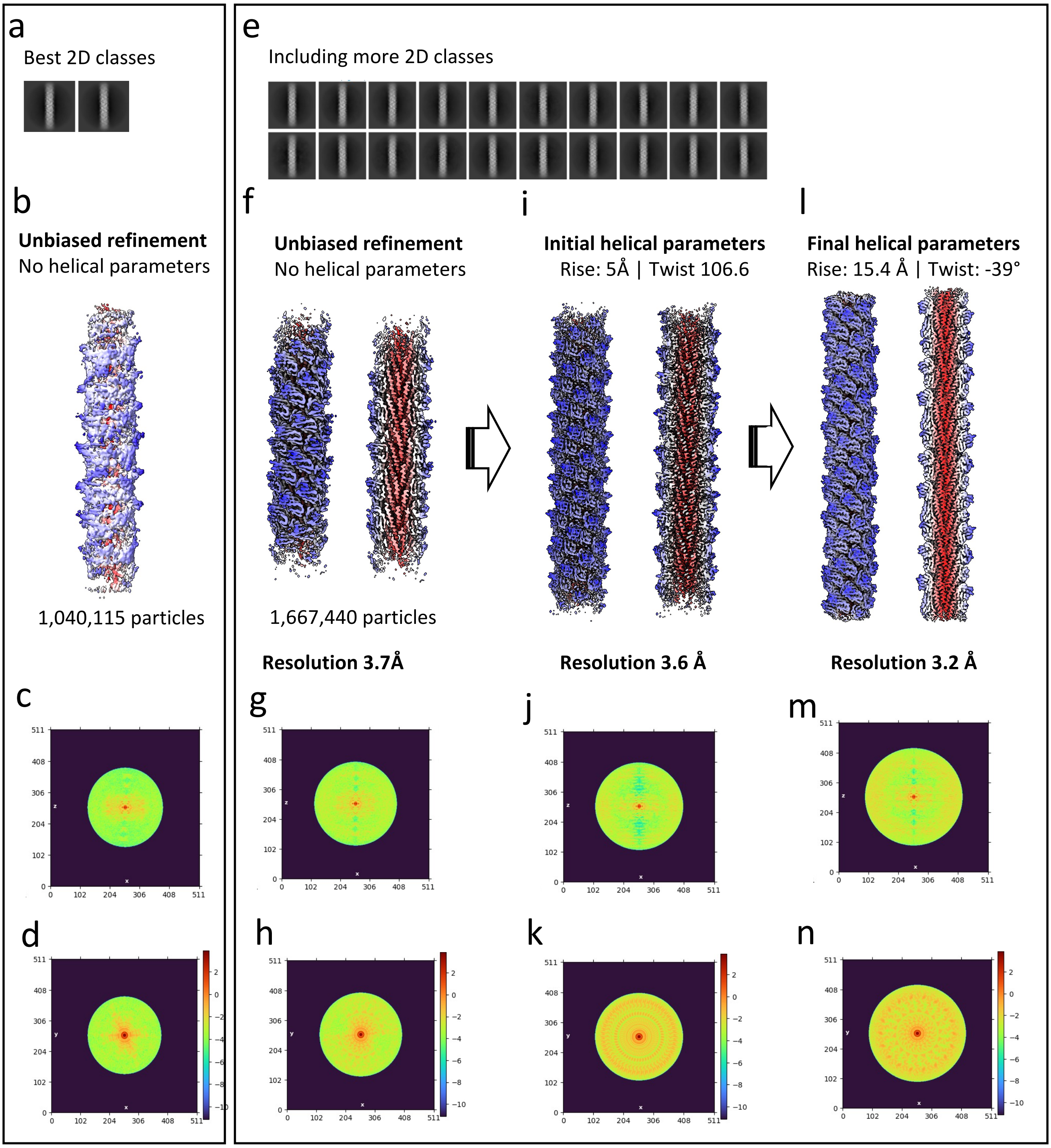

**Supplementary Figure 3 – Unbiased helical refinement of Aap**

**a**, the two best 2D classes used for the initial 3D refinement. **b**, 3D refinement without helical parameters resulted in a map with clear artifacts resulting from a limited number of views. This was also evident in the power spectra. While the power spectrum corresponding to the long axis of the filament was affected very little (**c**), the power spectrum relating to the short axis showed clear gaps in the signal (**d**). **e**, adding more 2D classes (including less well resolved ones), resulted in an improved 3D refinement (**f**) with 3.7 Å resolution. While there was still some missing information in the power spectrum (**h**), molecular details could be discerned, suggesting a helical rise of 5 Å and twist of 106.6°. **i**, Applying these parameters in a new round of helical refinement resulted in a map with nominally increased resolution (3.6 Å). However, further inspection and initial model building based on the unbiased map (**f**) showed that the archaellum was composed of subunits adopting three different conformations, revealing more accurate helical parameters of 15 Å rise and -39° twist. Applying these parameters to helical refinement improved the resolution further to 3.2 Å (**l**). Note that the power spectrum corresponding to the initial helical parameters (5 Å rise and 106.6° twist) lead to a scrambled power spectrum (**k**), while that of the final parameters (**n**) improved the signal pattern of the power spectrum of the unbiased map (**h**).

**Supplementary Figure 4**

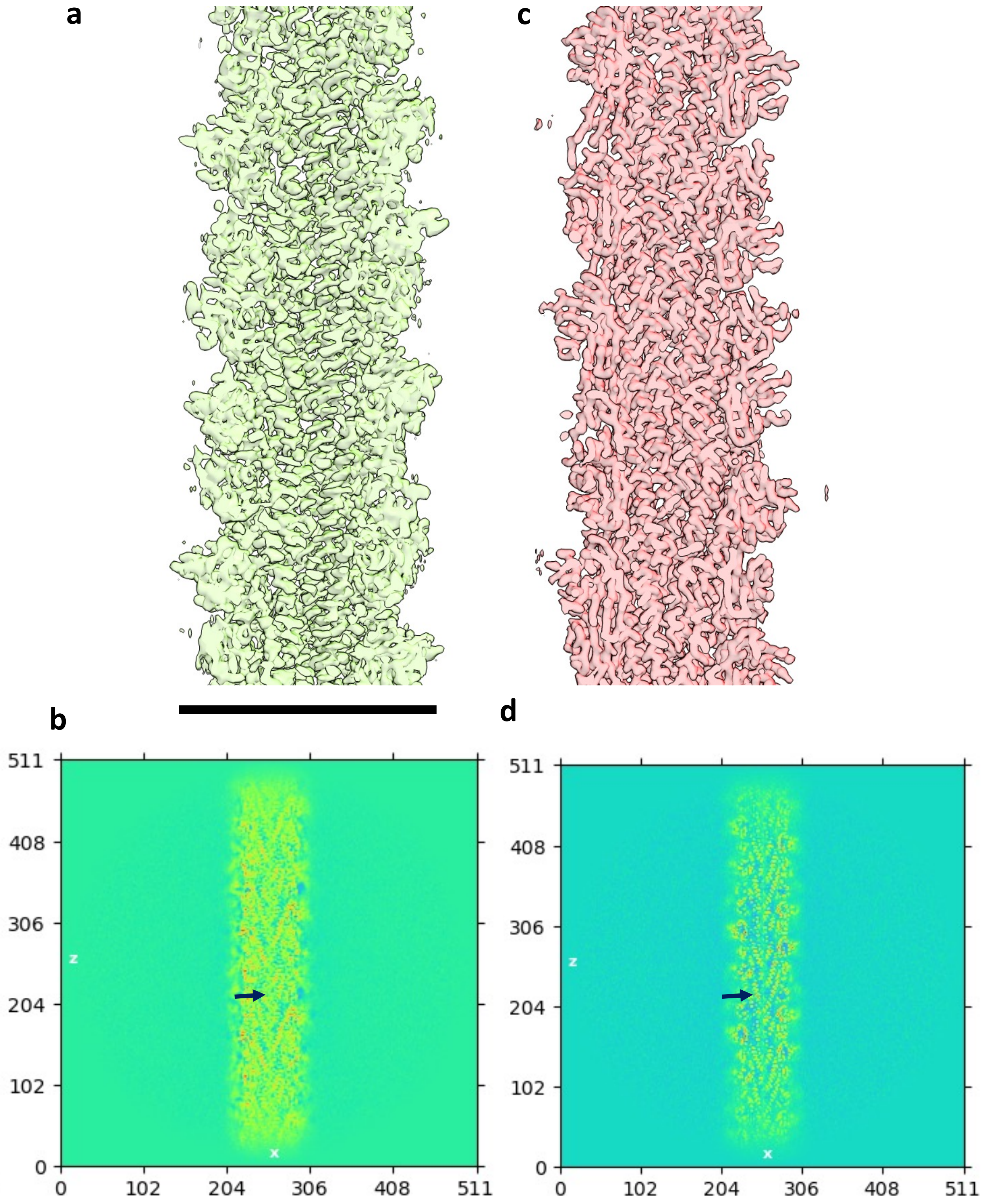

**Supplementary Figure 4 – Updating helical parameters improves the resolution of the map**

**a,c,** surface representation in cross-section (**a**) and projection (**c**) of the map obtained with 5 Å rise and 106.6° twist. **b, d**, surface representation in cross-section (**b**) and projection (**d**) of the map with map resulting from applying -39° twist and 15.4 Å rise. Note that the quality of the map, particularly the core is greatly improved with -39° twist and 15.4 Å rise. Scale bar 80 Å

#### Supplementary Figure 5

a

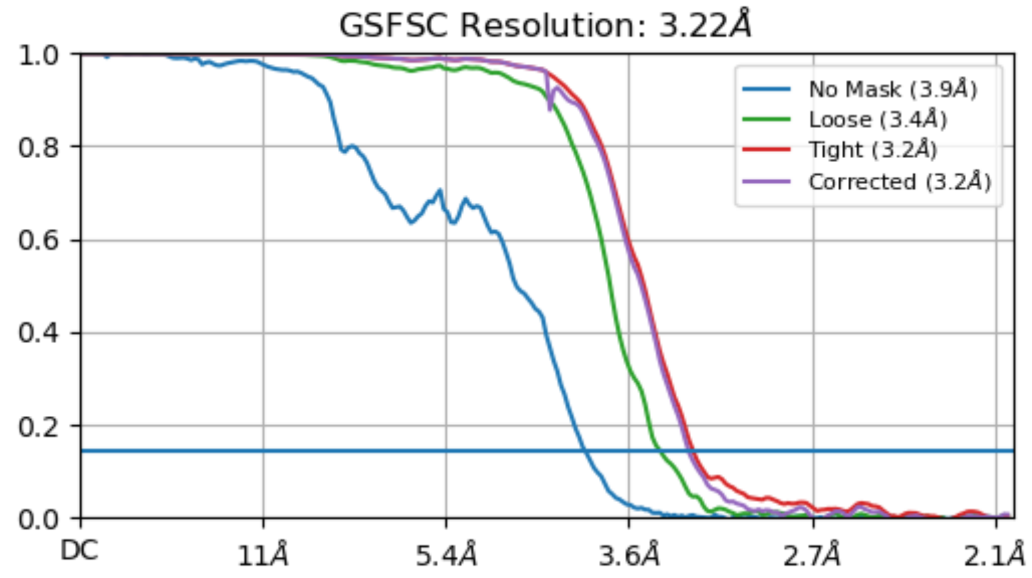


b

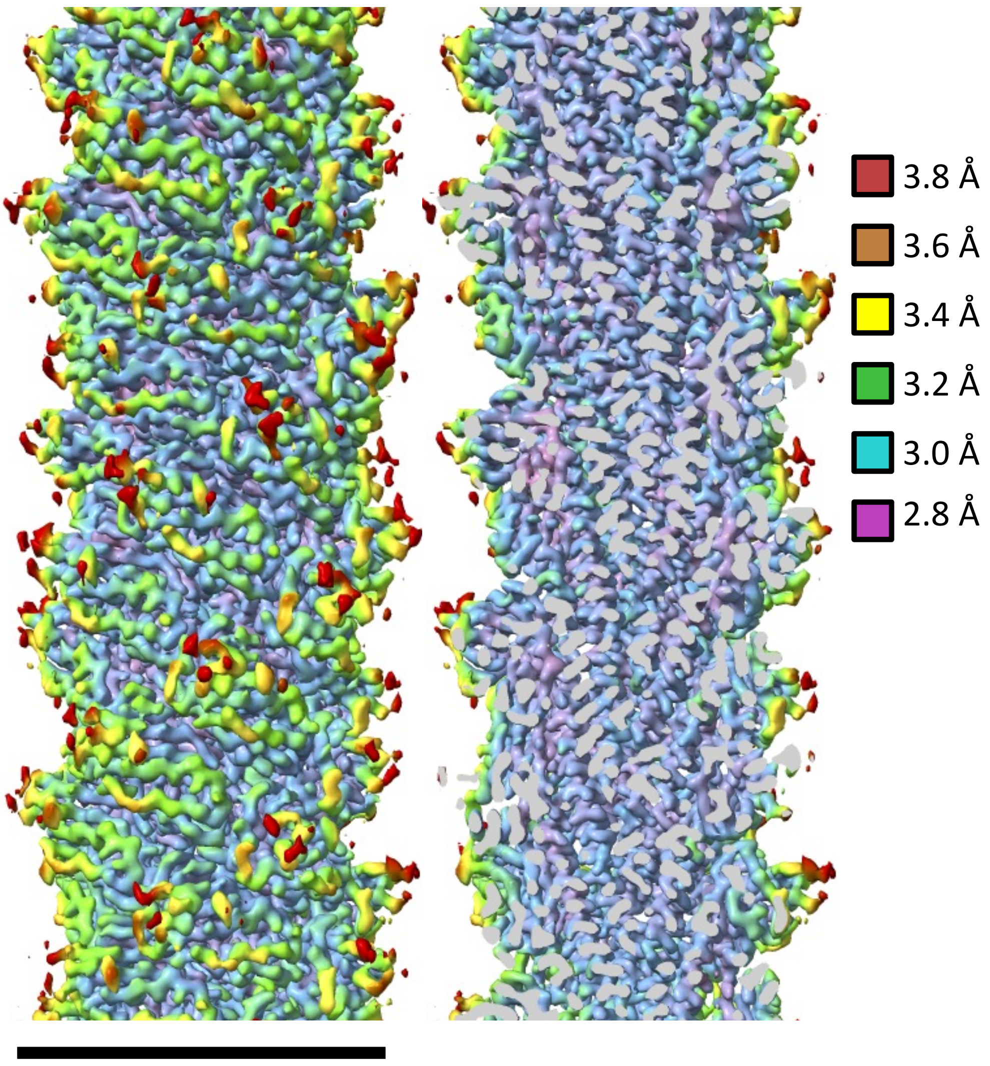

**Supplementary Figure 5 – Resolution estimation**

**a,** Fourier Shell Correlation (FSC) showing that the global resolution of our map is 3.2 Å. **b**, local resolution estimation suggests that the core of the filament reaches a resolution value of ~2.8 Å, while the periphery of the filament (including the surface glycans) is less well resolved (~3.8 Å). Scale bar 80 Å.

### Supplementary Figure 6

**a**

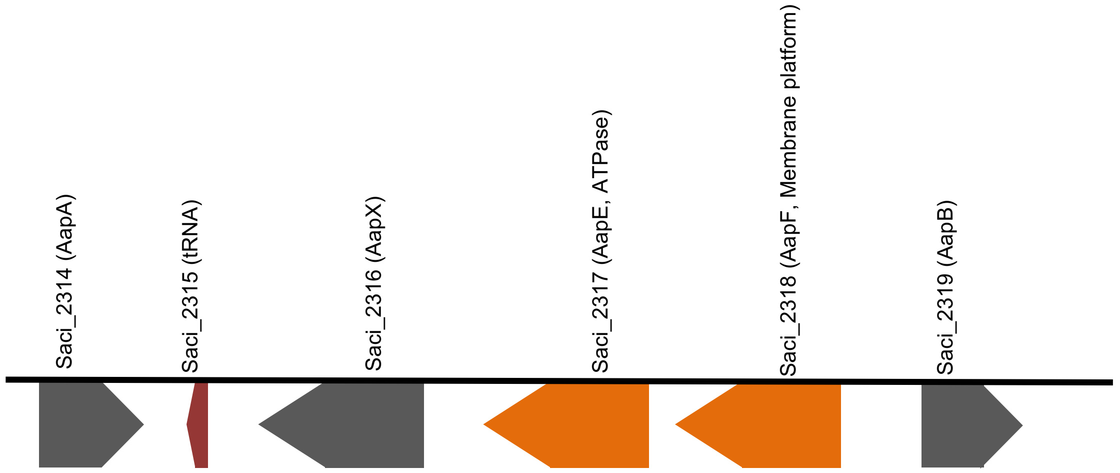


**b**

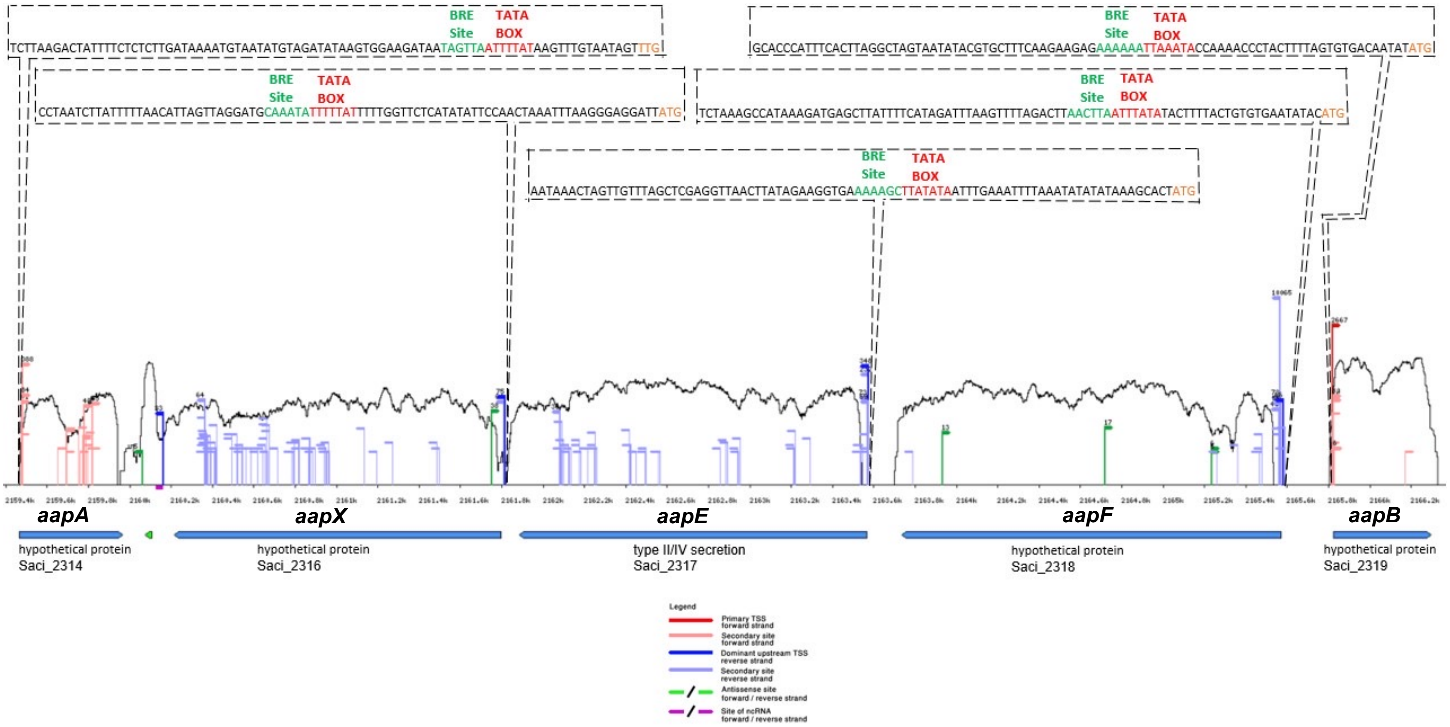

**Supplementary Figure 6 – Aap gene cluster and transcriptomics profile**
**a**, Schematic of the Aap gene cluster in *S. acidocaldarius*, encoding for the genes aapA, aapX, aapE, aapF and aapB. **b**, RNA-seq expression profiles for each gene of the aap cluster. BRE sites within promotor regions are highlighted in green, TATA boxes in red [1].

### Supplementary Figure 7

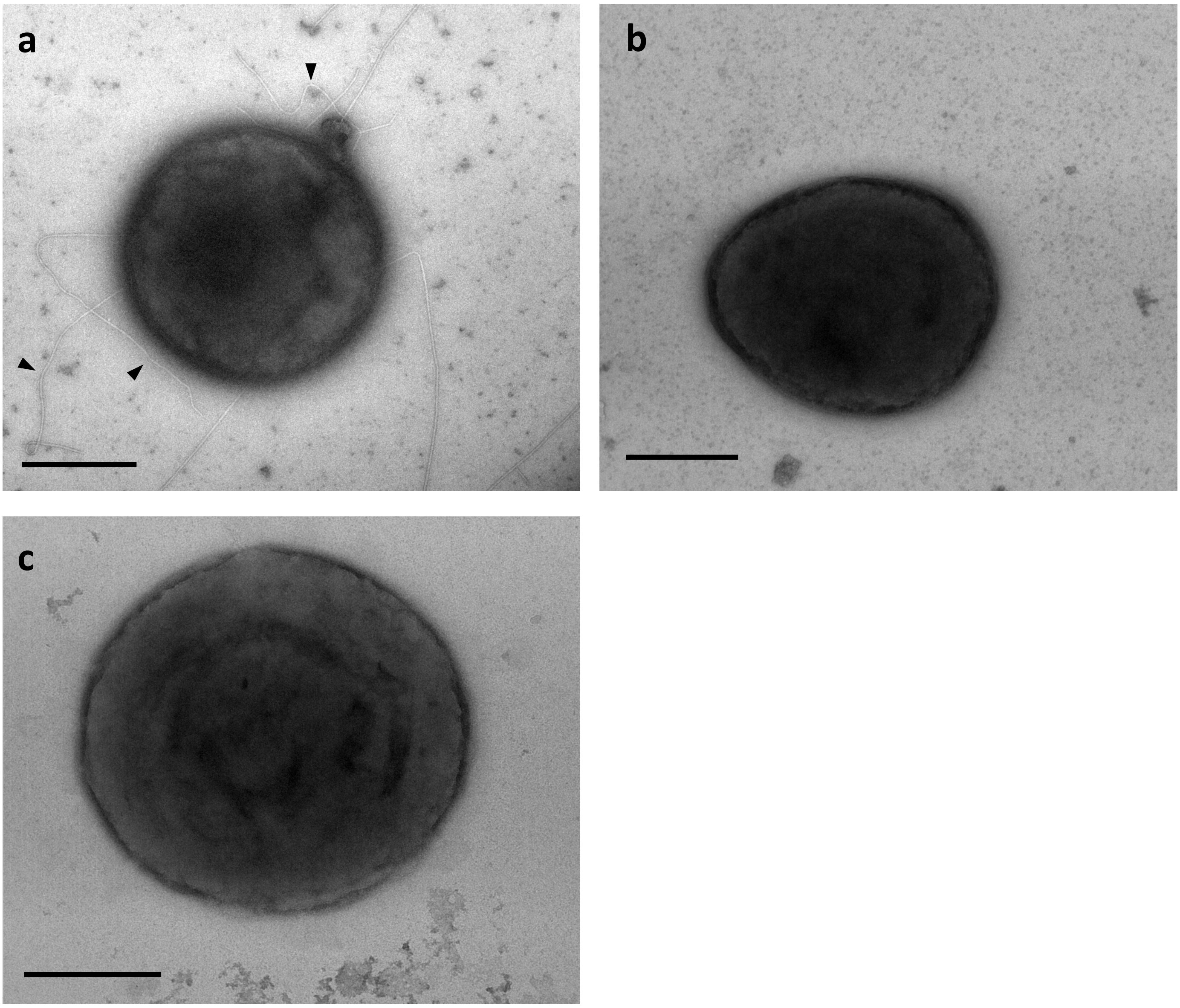

**Supplementary Figure 7 – Negative stain electron microscopy of *S. acidocaldarius* mutants**
**a**, the Δ*aapA* knockout strain still assembles AAP (arrowheads), while Δ*aapB* **(b)** and Δ*aapAB* double knockout (**c**) do not produce Aap.

#### Supplementary Figure 8

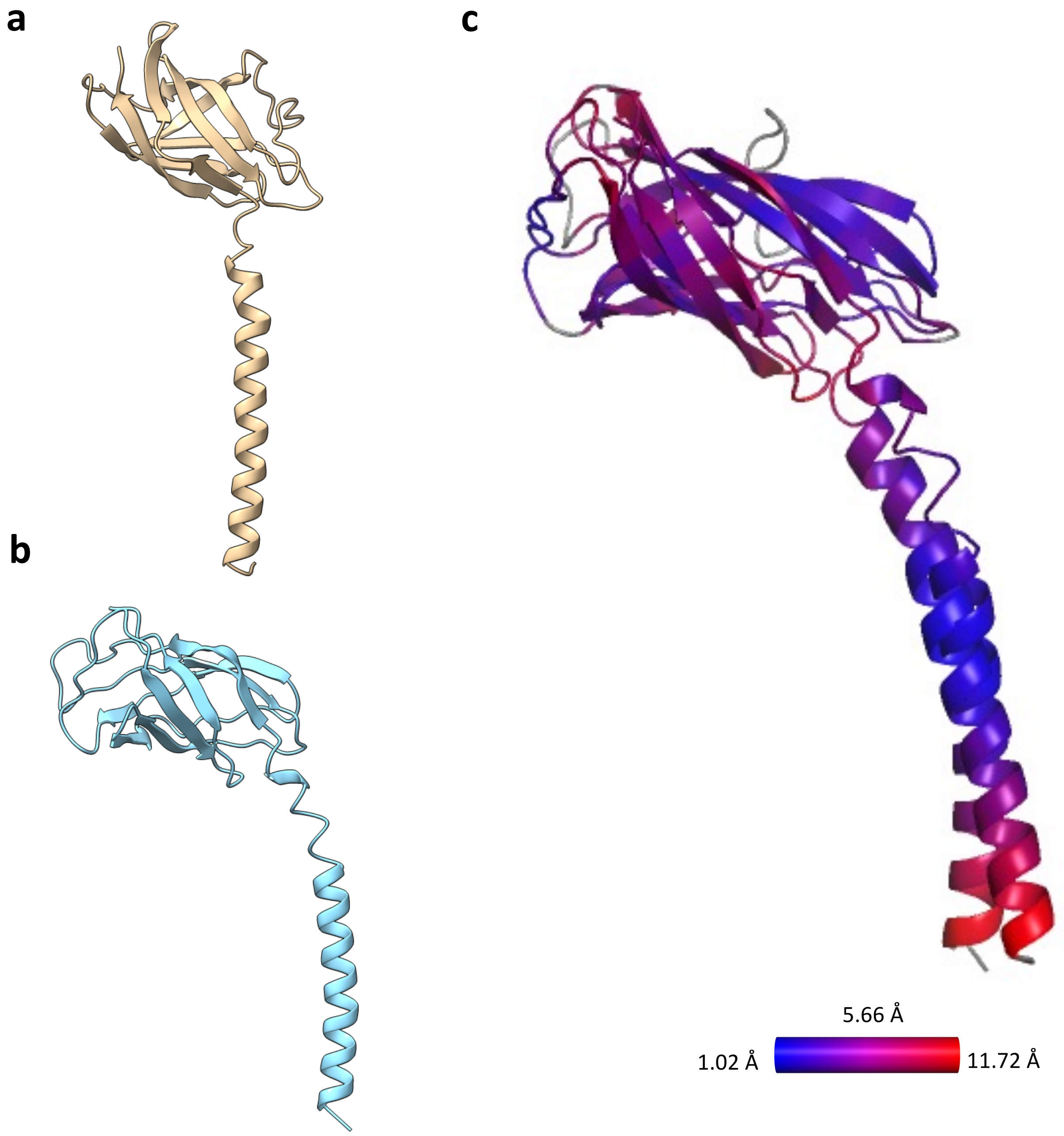


**d**

```
AapB      -----MNIEVKKSKKKNMRALSGAIVALILVIAGVIIAIAVVLFAFGLIPGISNQGSIQV  55
AapA      MYNKITMISRYRYDKRRIRALSGAIVALILVIAGVIIATAVVLFAFGLIPAISNQGSAQV  60
                 *.  : .*:.:******************** ***********.****** **

AapB      LGSGTITNSTASGSSRTIYNITITVKNTGT-TSISVTSININGQPFNING----------  104
AapA      VGTGAIEQAGS-----GQYNIIITVRNTASNFNVSVTSINIAGISFTINKINNITYNPNN  115
          :*:*:* :: :       *** ***:**.:  .:******* *  *.**

AapB      TAPSIPAGRTQPITFEVTPASGKPNFSPGASYTATIYFSNGQGAPATLIYQG  156
AapA      PMEQVGPGKTETLTITATPT-SSIVFSSGQTYTATVYFSNGLGAPTTLIYQG  166
             .:  *:*: :*: .**: ..  ** * :****:***** ***:******
```

**Supplementary Figure 8 – AapA and AapB from *S. acidocaldarius* in comparison**

**a,** AlphaFold2 prediction of AapA. **b,** experimentally solved structure AapB in A conformation. **c,** RMSD comparison between AapA and AapB showing their structural homology. **d**, sequence alignment between AapA and AapB. Black arrows show the N-glycosylation sites in AapB and a red arrow indicates a consensus (NXS/T) N-glycosylation sequon that does not appear to be glycosylated in AapB. The blue arrows highlight predicted (NXS/T) N-glycosylation sites in AapA. Scale bars 40 Å.

### Supplementary Figure 9

**a**

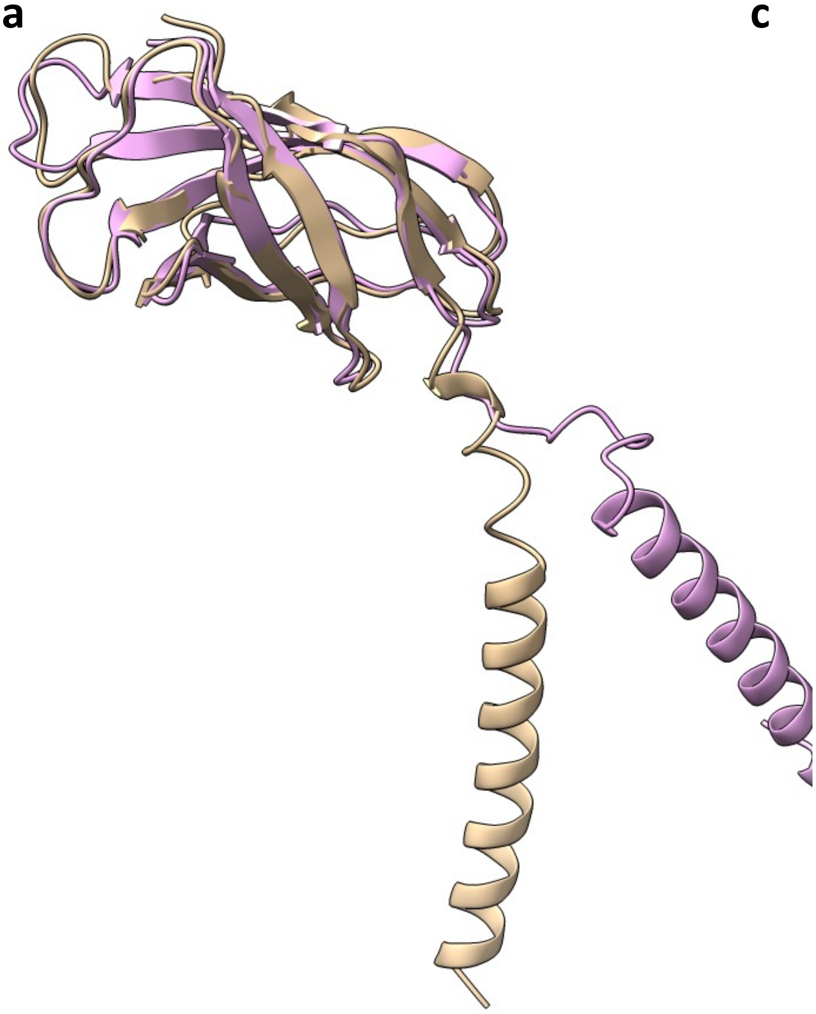

**c**

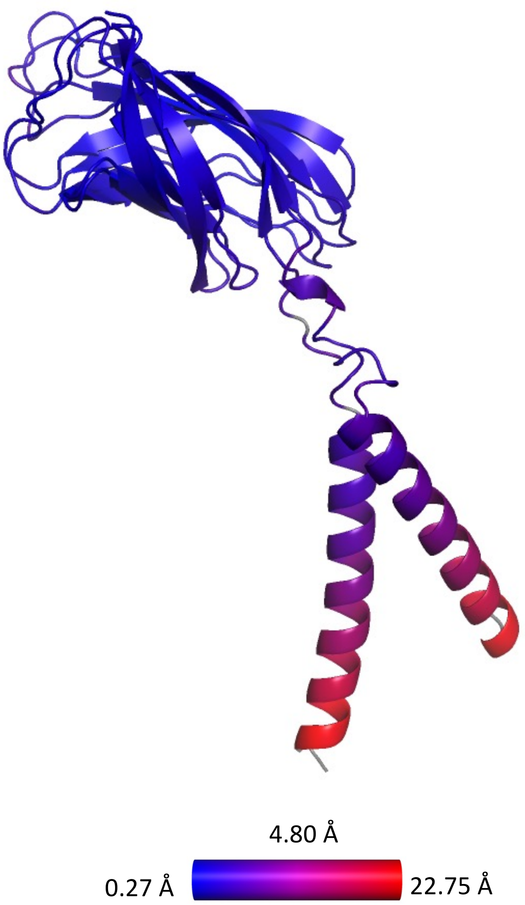


**b**

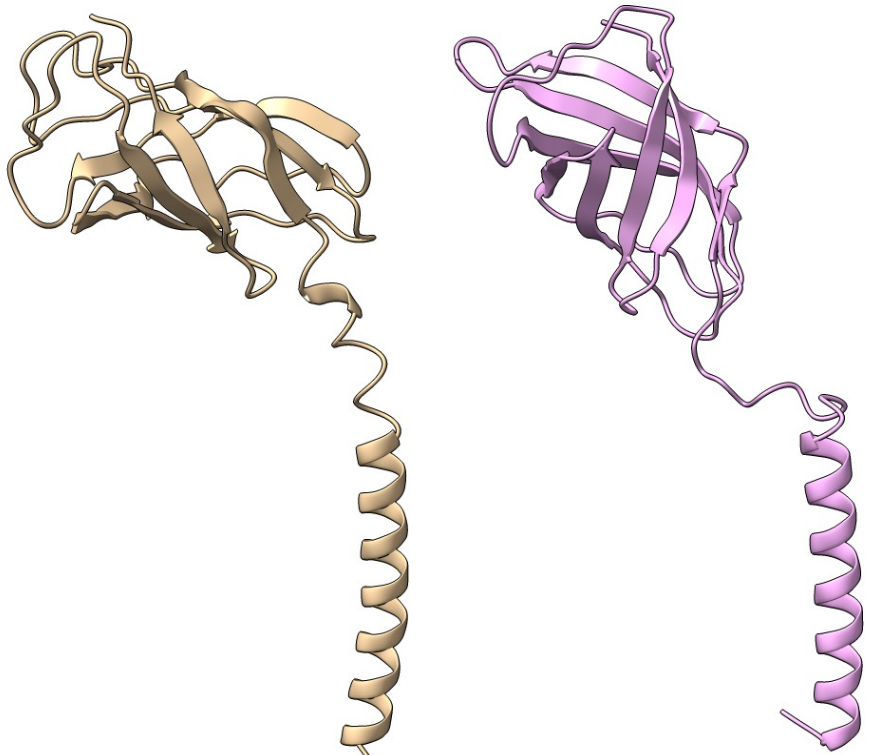

**Supplementary Figure 9 – Experimentally determined AapB structure vs. Alphafold 2 prediction**
**a,** superimposition of the experimentally solved (beige) and Alphafold-predicted (pink) structures of AapB. **b,** side by side comparison of the two structures from **a. c,** RMSD plot between the two structures from **a.**

#### Supplementary Figure 10

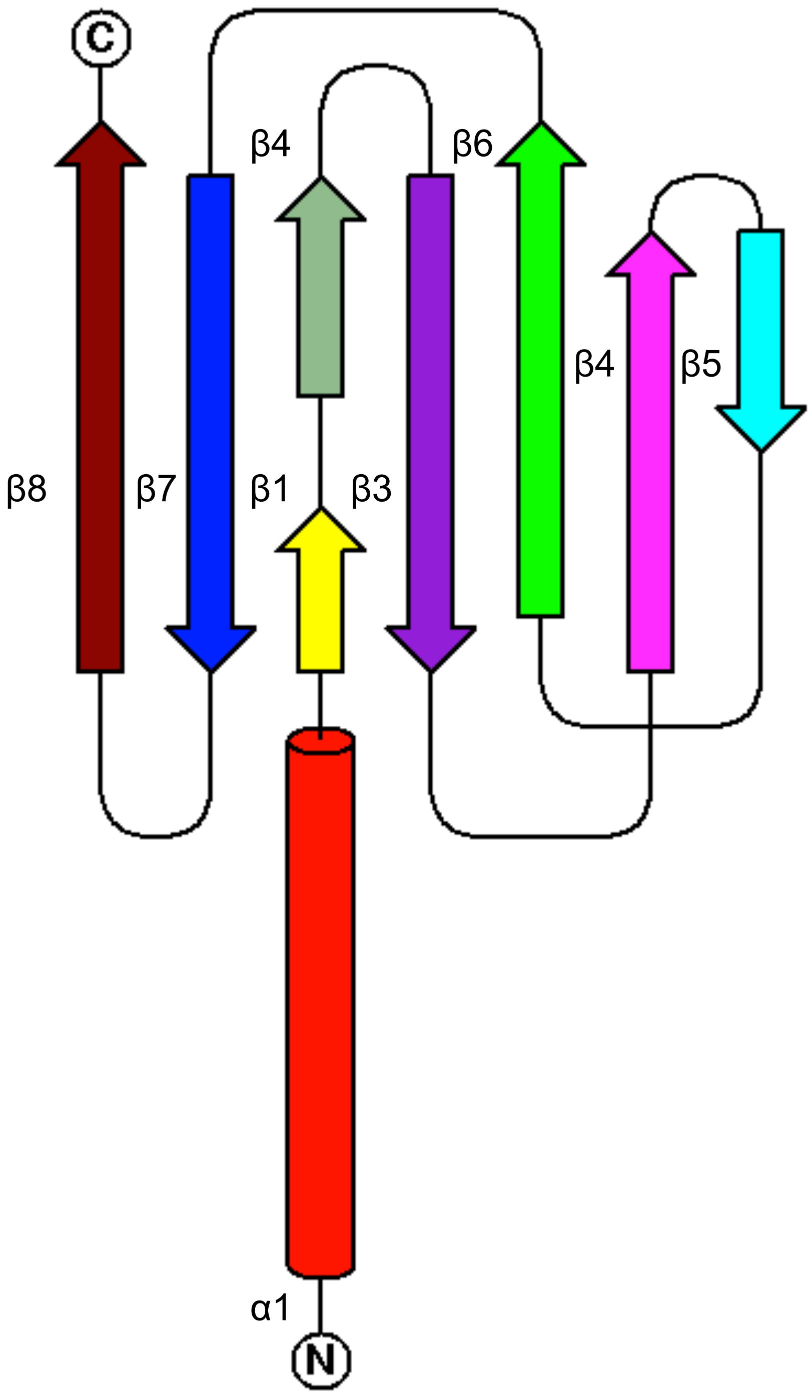

**Supplementary Figure 10 – Topology of AapB**

Topology diagram of AapB, showing α-helices as cylinders and β sheets as arrows.

**Supplementary Figure 11**

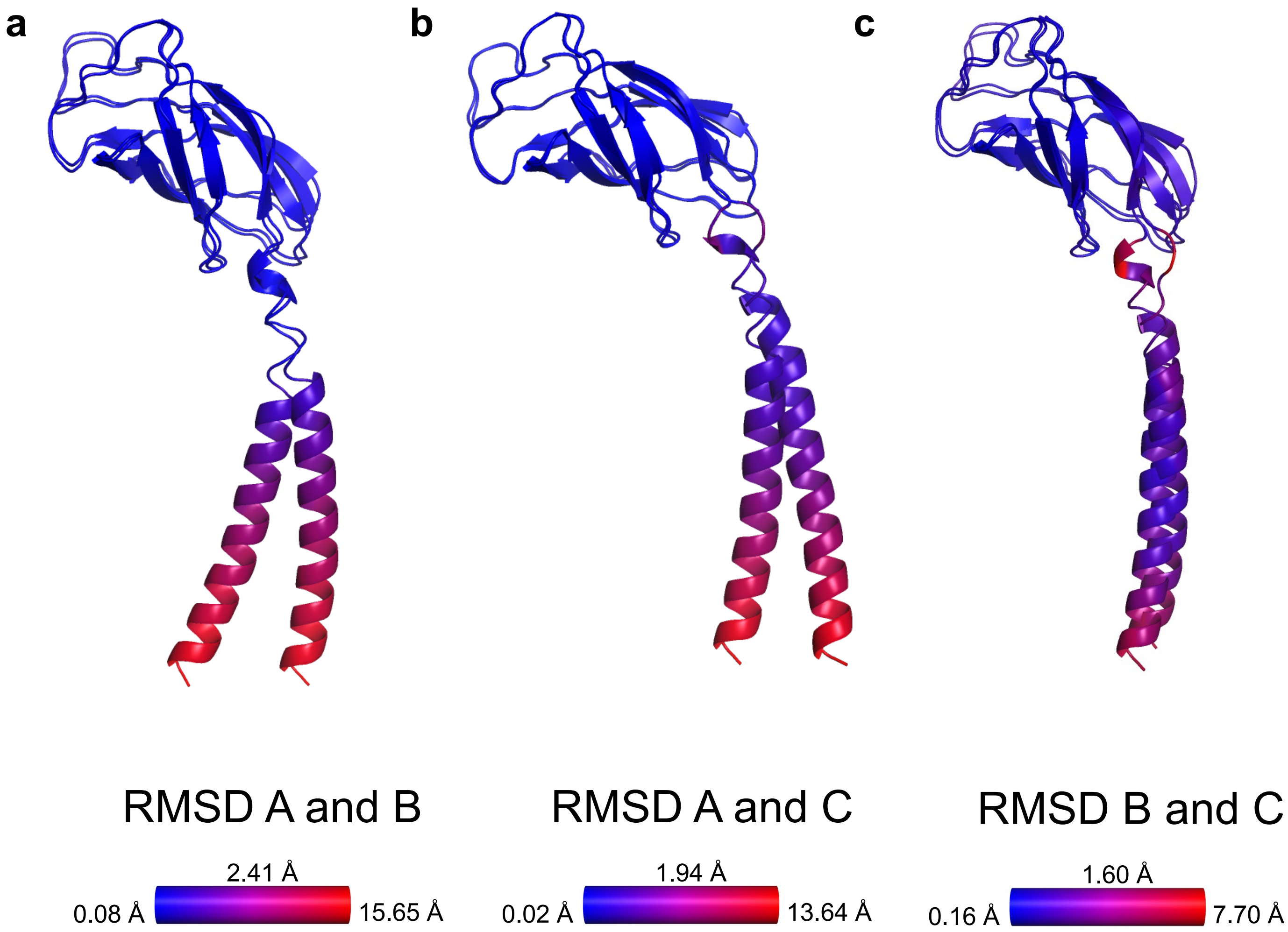

**Supplementary Figure 11 – RMSD between the three conformations of AapB**

RMSD between conformations A and B **(a)**, A and C, and B **(b),** and C **(c)**.

#### Supplementary Figure 12

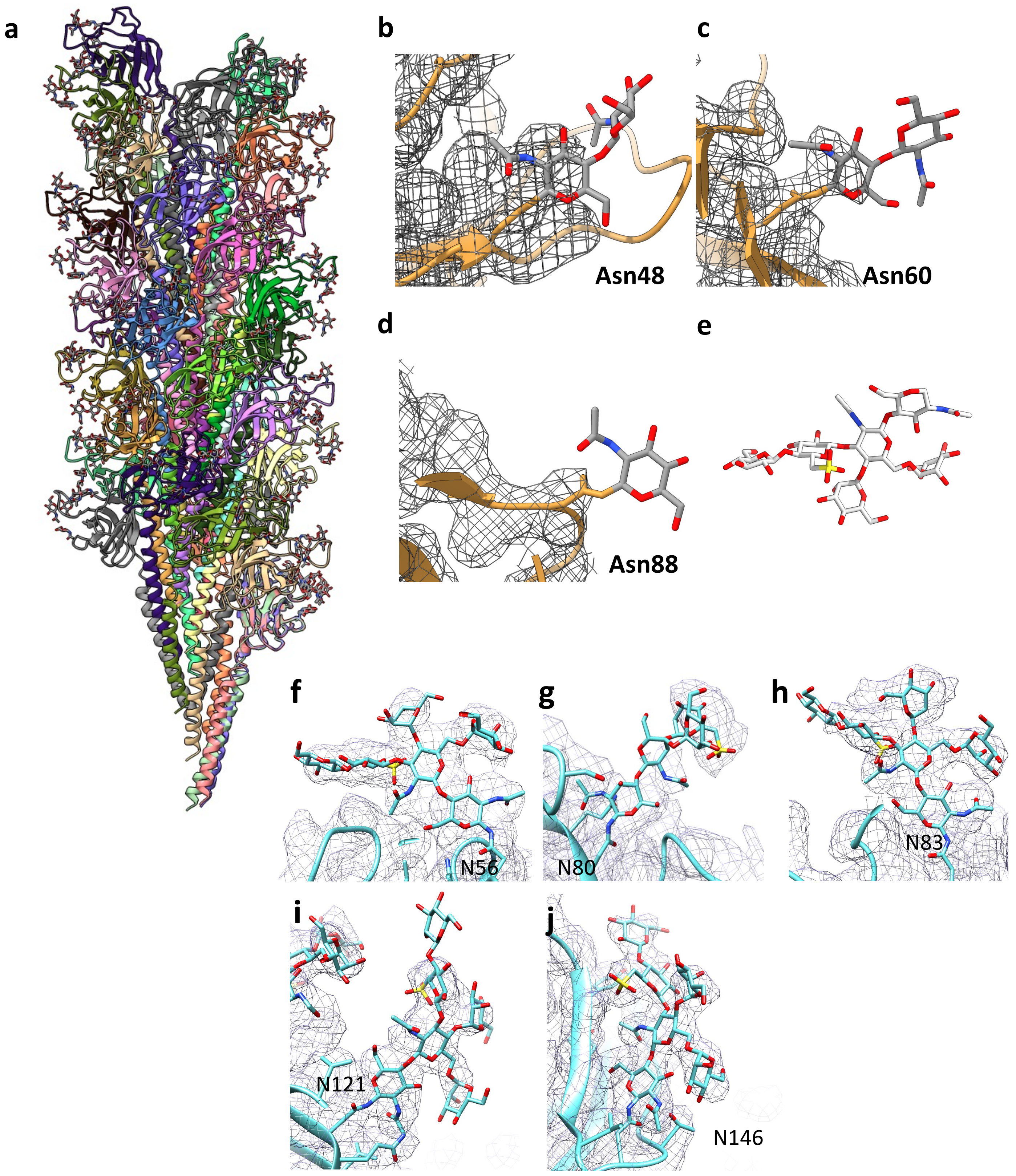

**Supplementary Figure 12 – Aap glycosylation**

**a,** atomic model of the *S. acidocaldarius* Aap showing the protein portion of the filament in ribbon, and the glycans in stick representation. **b-d,** three glycan models (beige) superimposed with the map (transparent grey) of AapB. **e,** structure of the complete *S. acidocaldarius* glycan. **f-j,** maps (grey mesh) and models (blue sticks) of the glycans found in the *S. acidocaldarius* thread filament.

### Supplementary Figure 13

**a**

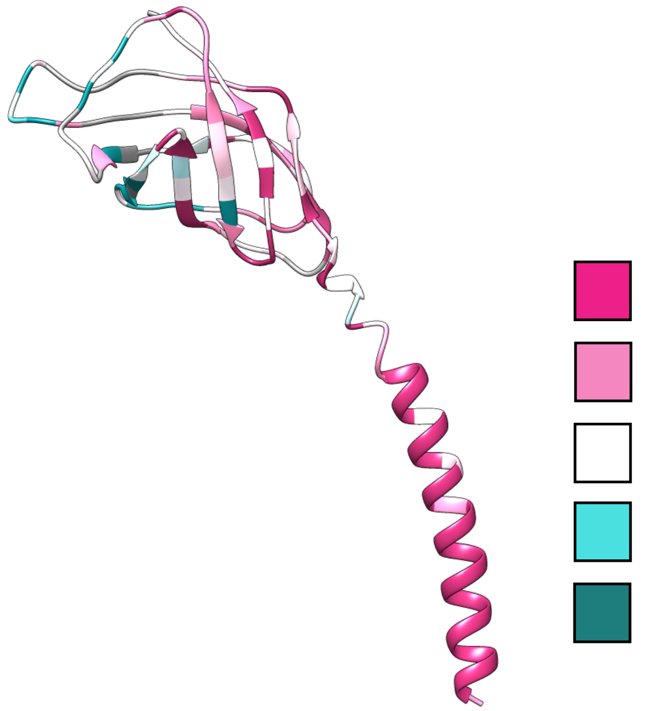


**b**

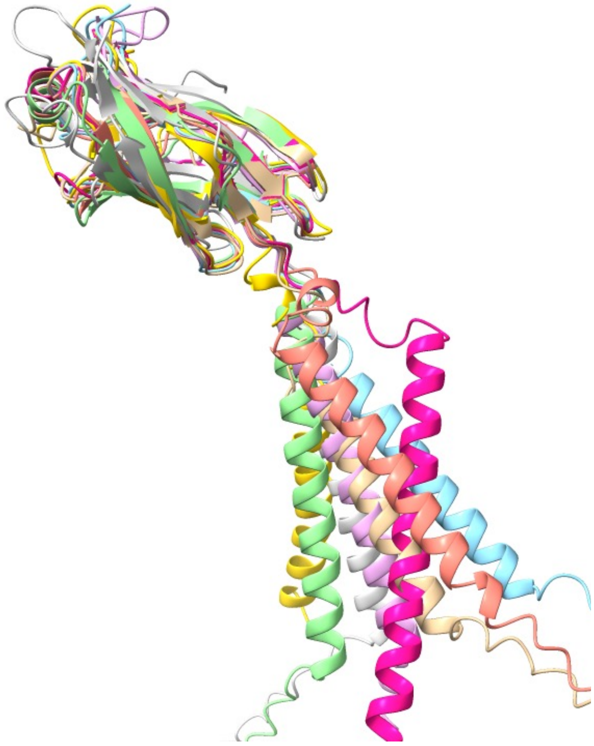

**c**

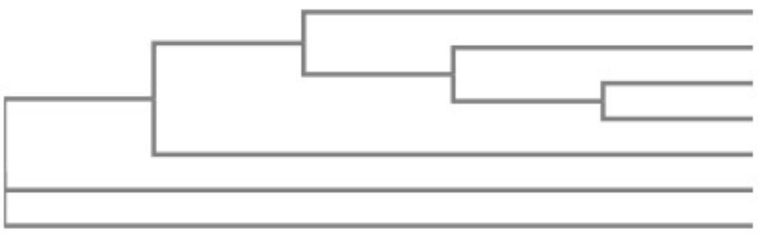


**d**

```
S.acidocaldarius    MNIEVKKSKKKNMRALSGAIVALILVIAGVIIAIAVVLFAFGLIPGISNQGSIQVLGSGT   60
A.brierleyi         --MKTINRKRRNQKGLSGAVTALILVIASVAIAIVVVGFVFGIFGAVTSTPSVTAAGTGS   58
S.shibatae          ---MKMVSRRKNAKALSGAVTALILVIASVIIALVVVGFAFGLFGAFSGQGTVSQVGTAT   57
S.islandicus        -----MISRRKNAKALSGAVTALILVIASVIIALVVVGFAFGLFGAFTGQGTVTQVGTAT   55
S.solfataricus      ---MDMASRRKNARGLSGAVTALILVIASVIIALVVVGFAFGLFGAFTGQGTVTQVGTAT   57
S.azoricus          --MSLKLKAKRSKKGISGAVTALILVIASVIIALVVVGFAFGLFGAIGGTPTVTQVGTGA   58
S.ohwakuensis       -----MLLRRKAKKGLSGAVTALILVIASVIIALVVVGFAFGLFGAFGGTPTVTQVGTGT   55
                             ::  :.:***:.*******.* **:.** *.**:: .. .  ::   *:.:

S.acidocaldarius    ITNSTASGSSRTIYNITITVKNTGTTSISVTSININGQPFNINGT---APSIPAG-RTQP   116
A.brierleyi         ISEIGN----TGQYTLTVVLDSTGSA--TVVSVSVDGYTTSV------STALSSGNNLIS   106
S.shibatae          LKT-SGT--------LIVTLKNTGANT-QVTGVLINGQSIPLTP----PYTISAGVTTYS   103
S.islandicus        LSASTGT--------LTVTLKNTGATT-QVTGAIINGNPVTLSS----QVTISAGQSTYS   102
S.solfataricus      LSASTLT--------LTVTLKNTGAST-QVTGVLINGNSGSVSG----MTTISAGVNTYT   104
S.azoricus          LYNSTSQ---KGYYEAIITLHSTGNV--QIVSATIVGTSIQVGTSDITGAALSAGANTVT   113
S.ohwakuensis       LTS---S---GNYVVATITLHSTGSV--QIVSATIAGTTYSATS---LSQSLSAGVNTVT   104
                    :                :.:..**     :..  : *             :: :*

S.acidocaldarius    ITFEVTPASGKPNFSPGASYTATIYFSNGQGAPATLIYQG--   156
A.brierleyi         ITLPTG-----FTPTPGQTYTIDLGLNDGQSVAVAAVYTTT-   142
S.shibatae          IALS-GSFTSSLSSLVGSTISLTLQLSNGQTVTVSAIVTS--   142
S.islandicus        ISLGS---SSTLQSLVGSTISLTLQLSNGQTVTVSAIITS--   139
S.solfataricus      ITISIGSISTTLRGLVGSTISLTLILSNGQTVTVSAIVTS--   144
S.azoricus          IYFPTA--AG-FTPQPGSVYQISLGLSDGTTVQVAVTYTGTT   152
S.ohwakuensis       LSFDTS--TGSVTLQPGSTYTISIGLSDGQTVQVAVVYQ---   141
                    : :             *      : :.:*  . .:
```

(Highlighted N-glycosylation sequons in S. acidocaldarius: NST, NIT, NGT, NFS; in S. azoricus: NST.)

**Supplementary Figure 13 – AAP homologues**

**a,** ConSurf comparison between *S. acidocaldarius* AapB and 6 homologs from related Crenoarchaeota. The model compares the conservation of each predicted structure, where pink indicates maximum conservation and green minimal. On the structural level, the alpha helix and inward-facing β-sheet of the head are the most conserved. **b,** superimposition of all structures shows the structural conservation between all 7 structures. Where no experimental structures were available, AlphaFold2 predictions were used. **c,** tree diagram showing the phylogenetic relationship between the seven homologs, based on the amino acid sequence. **d,** sequence alignment of all 7 homologs, where the yellow highlighted sequences are predicted N-glycosylation sites.

### Supplementary Figure 14

a b c d

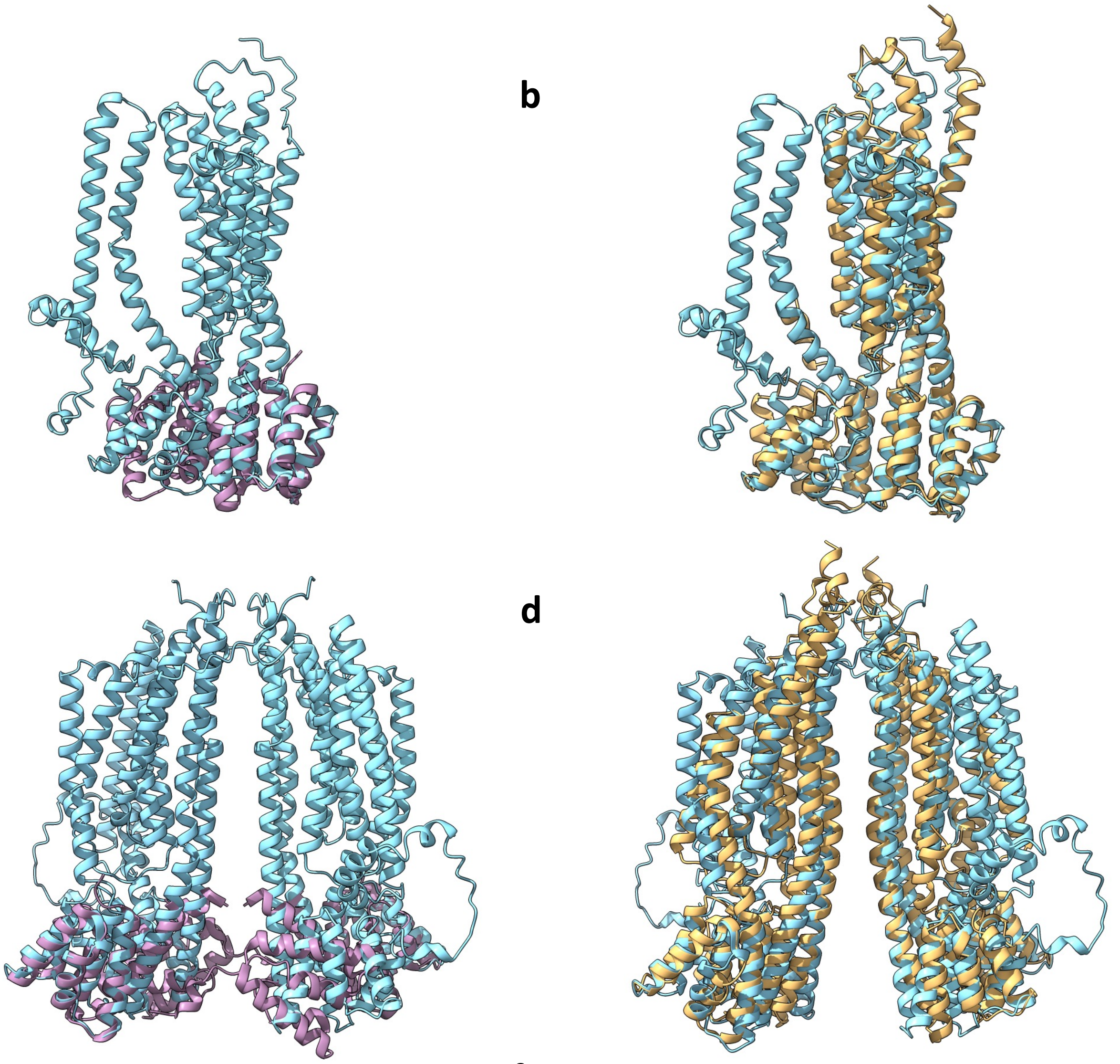

e

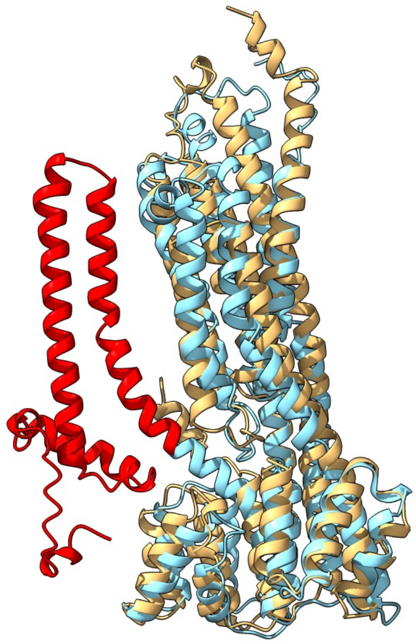

f

```
ArlJ           ------------------------------------------------------------   0
Saci_2318      MSRMSKDKKSSSNVNIPSIYLLFYHTPVVKRLAGYFDKRLLTSRNPEDPKLFASRLFLIL   60


ArlJ           -------------------------------------------------------MSVIF   5
Saci_2318      LVCIVLAVMFISFALIIFLRFYRVTLLPAYLALSLVMLFLGVIIPPIAYLISILDISQKI   120
                                                                      :*  :

ArlJ           DRNKKEEMDSKYIFMLAFMMALFSAGLPPETVLLKVSNKDSFHPYLKV----FRRIKNLV   61
Saci_2318      DKIKNGVDAESFSFA-TLFVIFLKSGLSPVLLFRKLEGSKAFSFVQDIVVYVNRRVQYLS   179
               *: *:    ..: *  :::: ::.:** *  :: *:....:*    .:     **:: *

ArlJ           SGFRYKFSQGINYAIRNSKVKFLSEFLVRLSQAVTFGDDMVQFLEREIDFSLAEYSSHSA   121
Saci_2318      E----SIEQALLKAMDINPGKLFNDFMLAYVTAIRTGAPVIETMEAKLKDLSKQFSLAAN   235
               .    .:.*.:  *:  .  *::.:*::    *:  *  ::: :* ::.    ::*  :

ArlJ           RVIESMNNF---LTVYATLNSSLAFLVADMTVLSVLYSGGSSILLQIFFLSTVILVNLTV   178
Saci_2318      LASDRLQGVAESYVVWLSSGYIMLYLVLIMGAI-LPFVGNSSSLLTVLGPVVVLVVP-MV   293
                . : ::..    .*: : .  : :**  *  .: : : *.** ** ::   .*::*   *

ArlJ           IMYYIYKPESY-MRYSSRDRLLLTALILLGIALNLTYTSFITMII---------------   222
Saci_2318      NLLFVYMADSLQLKFPETQSSA-YKIFYISLPIGLVV-AFLIMIFEHQIVYFITLSGGLQ   351
                : ::*  :*  ::: . :       :: :.: :.*.  :*: **:

ArlJ           ----------TGVLYLGVG--MRYRIFENKINNLERYFLLFTRYFTRNYSVVNNLKESLM   270
Saci_2318      NVFPVSIALLIGLLIASAPPAFFYQKEMREKSGFEEYAVKF----------LNAISEGLL   401
                          *:*  ..   : *:   .: ..:*.* : *          :* :.*.*:

ArlJ           AVL-----------RGDLGSAKPMIRRALNRLNLGVNKSKIFSLMGRESKSVLVTMLSEI   319
Saci_2318      AGLTFESIVTRLKDAQEMGKFREVLRKVDGYLKLGYPLTIAL-KRGADSIN---EFTSRI   457
               * *            ::*. : ::*:. . *:**   :  :   * :* .    : *.*

ArlJ           LYETISNGGNMLITGEIL----SKIGD-VVLNIRARKEQNGRAFEASIYALQVSSAGVSA   374
Saci_2318      ALYTL---SDMIEIGSMTPDNVRALADQINSQLVVRREYQGKVKPL----IATPYAGVLV   510
                  *:   .:*:  *.:        :.* :  :: .*:* :*:.        : .  *** .

ArlJ           ALISITSMLNNLFSIGELASVIAFNPIDIGFVSKIFLIMLFV-----MSFANGMAI-SLA   428
Saci_2318      SLIATFLLASGILSM-LNSGIAVYGPIATGLVSIPQIIFITAISGILNAFLAGLLIGKIG   569
               :**:   : ..::*:   :.: .:.**  *:**   :*:: .      :*  *: * .:.

ArlJ           YGKSIYASLYFIGL-LMIMSAISYYVILTLTAQLFQAFSSGGIISGIQNTT   478
Saci_2318      YGKASAG--FIHGIILMIVVTLTIFAFVELRISIVPNFHSNISF-------   611
               ***:  .  :: *: ***: ::: :.:: *  .:.  * *.  :
```

**Supplementary Figure 14 – Alphafold prediction of AapF compared to ArlJ, and PilC**

**a,** Alphafold2 prediction of an AapF monomer (blue) superimposed with the structure of the bacterial PilC from *Thermus thermophilus* (PDB-2WHN,[4]). **b,** Alphafold2 prediction of an AapF monomer (blue) superimposed with the AlphaFold2 model of *S. acidocaldarius* ArlJ (beige). **c,d,** dimer models of **a,b**. **e,** comparison between the Alphafold models of AapF (blue) and ArlJ (beige). The extra N-domain of AapF has been highlighted in red. **f,** sequence alignment between ArlJ and AapF.

#### Supplementary Figure 15

**a**

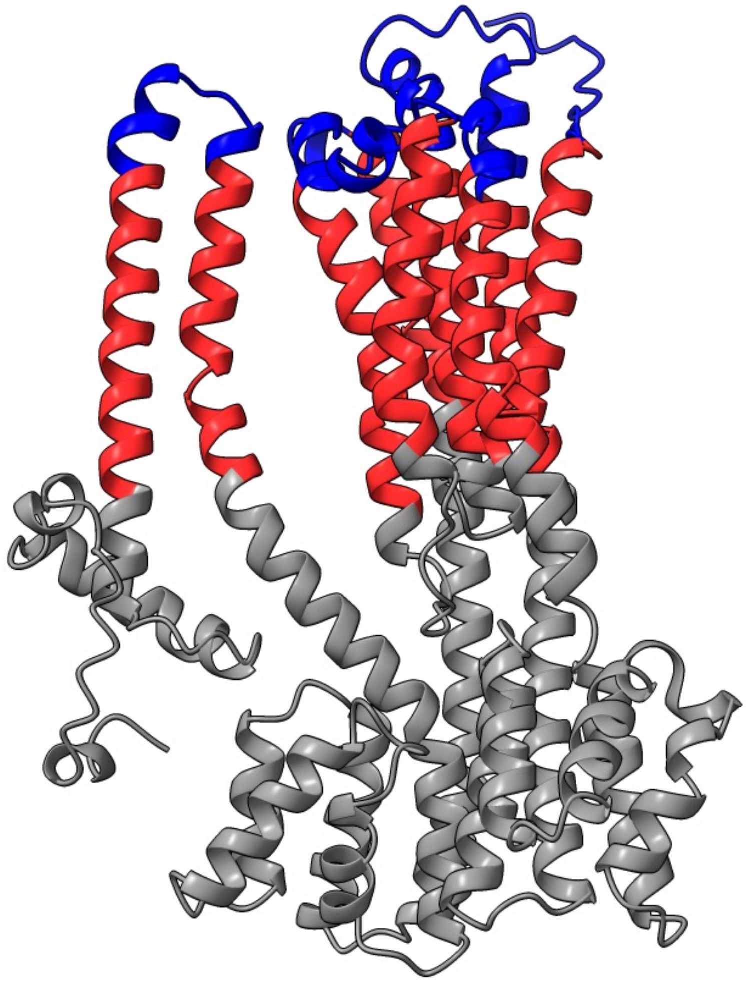

**b**

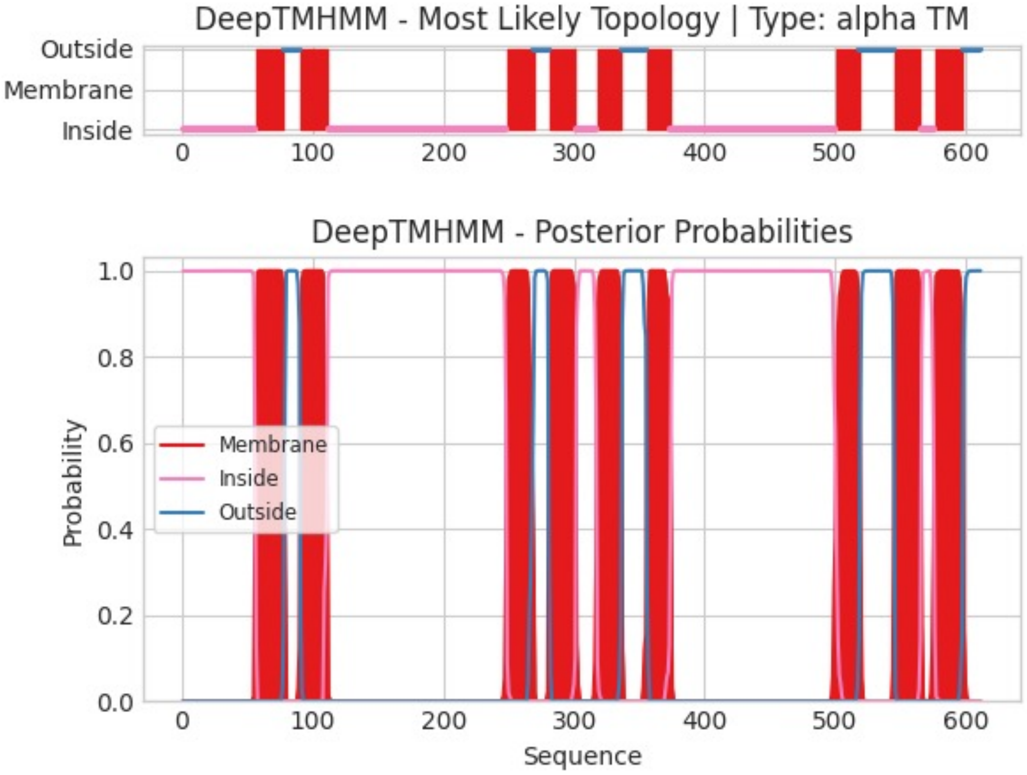

**Supplementary Figure 15 – DeepTMHMM topology prediction for *S. acidocaldarius* AapF**

**a,** colour coded topology prediction for AapF. Blue, outside; red, membrane integral; grey inside. **b,** probability plots for the predicted topology.

### Supplementary Figure 16

**a**

```
ArlI        -----------------------------------MSFVEDYLTKL-QERPTIIENPNI   23
Saci_2317   MALSKFTRKSNKDLEEVSQYKIPATLYPLTGIPEEVLNISSEYEVDILSQIPKEL--VDQ   58
                                   :.: .:* ..: .: *. :   :

ArlI        LKGSKIFNAIYRVDDFVYIHIQSIKSEDGYNQYNVIEPPRPTHDEME------EIEEKFA   77
Saci_2317   LNNSGIEMKIPNPHVFIY-----FDNDKGVHKYVLVEPPLEEKSFVIYKLLIKYLERQFV   113
            *:.* *   * . . *:*    :...:* ::* ::***   :. :       :*.:*.

ArlI        ---LSIGDKEPPEDTKEKEKLIRSILDKILLRMRLSVPKEYVIYHFIRDKLYTGSLEPLI   134
Saci_2317   GKHFDIGASIRDFSNIYRDYVIEGSRG---DYTVLKPEARVALYHIIRNLLGYNVFTPLL   170
              :.** .   ..  :: :*..  .      *.   . ..:**:**: *   . : **:

ArlI        RDPYIEDISIPGLGH-VYIVHKVFGPMRTSIKFENY-----------EELDNLIVSLSEK   182
Saci_2317   ADYKVEDISVNGLNVPIYVYHRDYEYIPTNIIFQKRMSVLNEEVDGEEFLNQITMRYISL   230
            *  :****: **.  :*: *: :  : *.* *::          * *::: :   .

ArlI        SYRPVSHNRPVVDASLPDGSRVNFVYGVDISRRGSNLTVRKFSRVPTSITQLIMFGTLSS   242
Saci_2317   SGRSISIASPISDGMLPAGDRIAATYGREVSAKGTSFVIRRFSESPITVLHLIKQGVISA   290
            * * :*   *: *. ** *.*:  .** ::* :*:..:.*:**. * :: :**  *.:*:

ArlI        MMAAYIWTMLDEGMNLFVCGETASGKTTTLNAITAFIPPNLKIVTIEDTPELTVPHSNWV   302
Saci_2317   DLVAYLWYAIDLKMSFMVIGTTGAGKTTVLNALLNLVKDTMKIVSIEDIPELKLARDNWV   350
             :.**:*  :*  *..:* * * *.:****.***:  ::  ..:***:*** ***.: :.***

ArlI        AEVTRETGGE--GTIKLFDLLKAALRQRPNYILVGEIRDKEGNVAFQAMQTGHSVMATFH   360
Saci_2317   QLYTRPVFGALGKEITLMDLLKLSLRYRPDIITVGEIRGEESYVLFQALSTGHGGATTFH   410
              ** . *     *.*:**** :** **: * *****..:*. * ***:.***.  :***

ArlI        AANITTLIQRLTGYPIEVPKSYINNLNIALFQTALYDKKGNLIRRVVEVDEIIDIDPVTN   420
Saci_2317   AYDPESAIKRLMNEPLNIPRDWIPMMNVVLTVRRI-PFGLSFKRRVVEVDEVVSYNKLKK   469
            * :  : *:** . *:::*:..:*  :*:.*     :     .: ********:: : :.:

ArlI        DVVYIPAFTYDSVQDKMLFAGKGSSYLIENKIAVKRGID--RRNIGLLYDELQMRSRFLN   478
Saci_2317   VGSWDPN------NDSHTFDLSKSEV-------LKSRIEEIGLNLDIVGDEIKRRANYLK   516
              : *      :*. *  .*.       :*  *:    *:..: **:: *:..*:

ArlI        LLVEKK-IFNYYDVWDYILR---ARQMGLEEAIKYVSNI-----   513
Saci_2317   MLSDIKQILDRPDAYVFVKKYIVLYSIKPDEAIKEVQRMSVAKV   560
            :* : * *::  *.: :: :    .:  :**** *..:
```

**b**

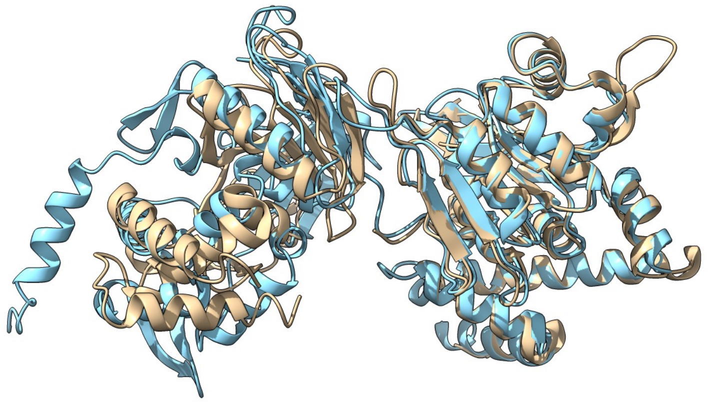

**c**

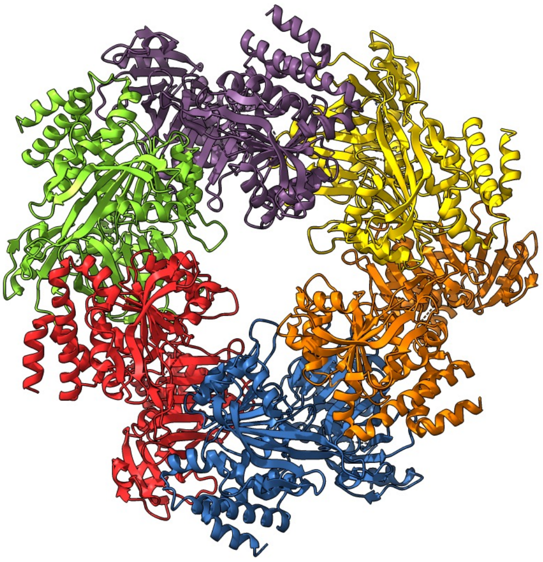

**d**

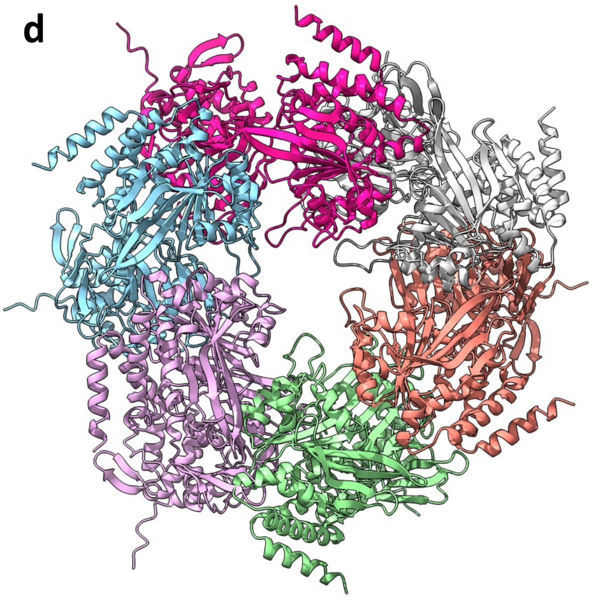

**Supplementary Figure 16 – Alphafold prediction of AapE compared with ArlI**

**a,** sequence alignment between *S. acidocaldarius* AapE and ArlI. **b,** superimposition of AlphaFold2 predicted AapE and the experimentally solved structure FlaI from *Sulfolobus acidocaldarius* (PDB-4II7,[5]). **c,d,** hexameric models of FlaI (left) and AapE (right).

#### Supplementary Figure 17

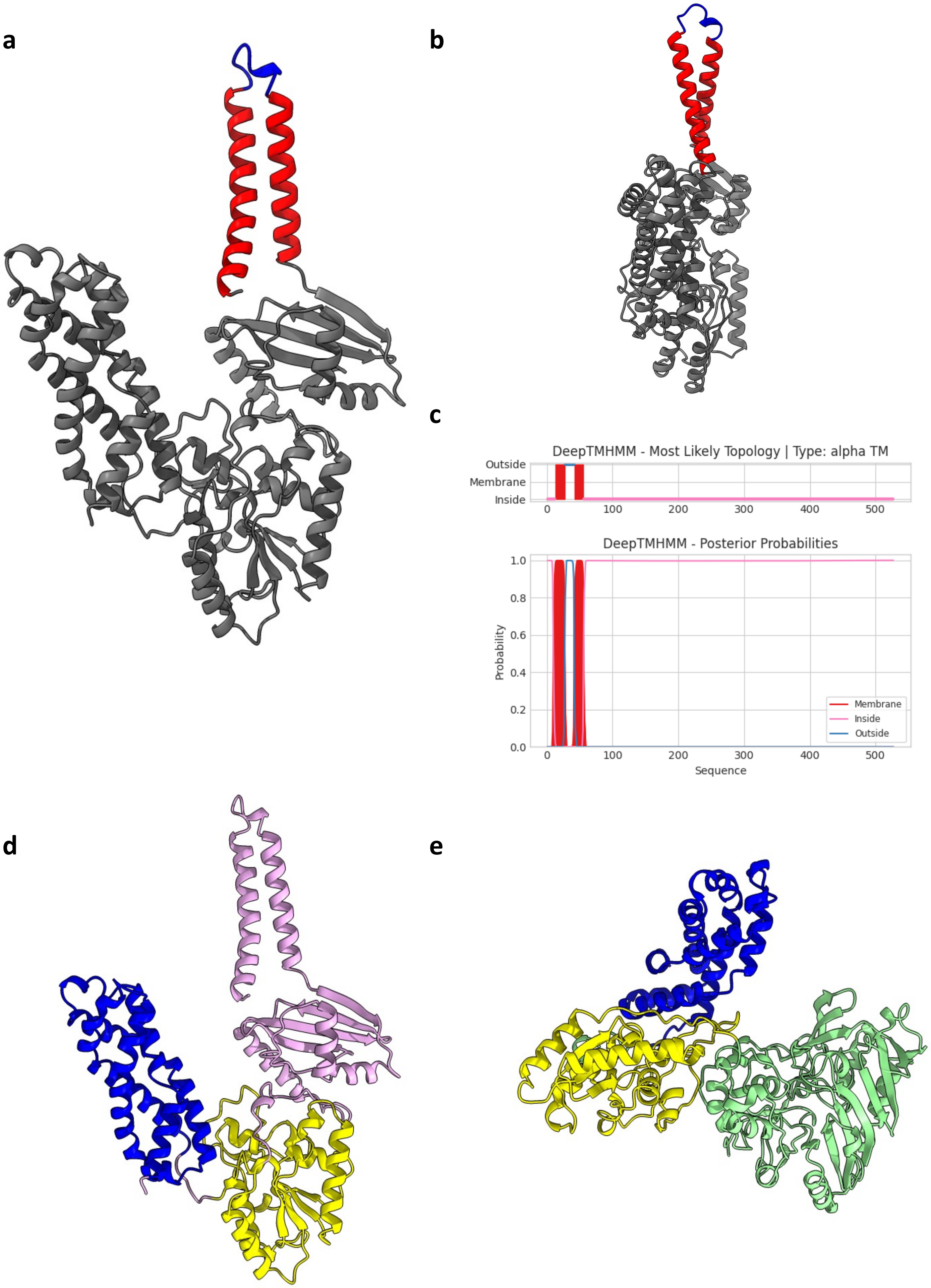

**Supplementary Figure 17 – Alphafold prediction of AapX**

**a,b** Alphafold model of the AapX monomer colour-coded for topology its topology, as predicted by DeepTHMHH. Blue, red and grey represent outside, membranal and inside respectively. **c,** DeepTHMHH probability plots for the predicted topology in **a, b. d,** the three distinct domains of AapE in blue, yellow and pink. Comparing *S. acidocaldarius* AapE with homologues, e.g. **d,** the FAD binding protein of *Sulfolobus solfataricus*,[3]. **e**, shows that in *S. acidocaldarius* AapE, the FAD domain (green) is exchanged for a membrane-binding domain (pink). Scale bar 50 Å.

### Table 1 - Cryo-EM data collection, refinement and validation statistics

| | **AAP**<br>**(EMDB-)**<br>**(PDB )** |
|---|---|
| **Data collection and processing** | |
| Magnification | 105k |
| Voltage (kV) | 300 |
| Electron exposure (e–/Å$^2$) | 42.33 |
| Defocus range (μm) | -2.0 to -3.5 |
| Pixel size (Å) | 1.047 |
| Symmetry imposed | Twist: -39.859° Rise: 15.403 |
| Initial particle images (no.) | 2,207,193 |
| Final particle images (no.) | 947,729 |
| Map resolution (Å)<br>FSC threshold | 3.22<br>0.143 |
| Map resolution range (Å) | 3.8-2.8 |
| Resolution (Å) (map/model; FSC = 0.5) | 3.22 |
| **Refinement** | |
| Initial model used (PDB code) | Ab initio |
| Model resolution (FSC = 0.50/0.143 Å) | |
| Model refinement resolution (Å) | 3.22 |
| Map sharpening B factor (Å$^2$) | 0 |
| Model composition<br>Non-hydrogen atoms<br>Protein residues<br>Glycans | <br>72,072<br>10,152<br>408 |
| B factors (Å$^2$)<br>Protein<br>Glycan* | <br>64.8<br>205.6 |
| R.m.s. deviations<br>Bond lengths (Å)<br>Bond angles (°) | <br>0.010<br>1.611 |
| Validation<br>MolProbity score<br>Clashscore<br>Poor rotamers (%) | <br>1.43<br>2.70<br>3.08 |
| Ramachandran plot<br>Favored (%)<br>Allowed (%)<br>Disallowed (%) | <br>99.0<br>1.0<br>0.0 |

*The refinement was conducted against the DeepEMhanced map, which blurred the glycan density resulting in elevated B-factors for those.
